# Semaglutide quiets “food noise” in mice by impairing the wanting, but not the liking, of glucose

**DOI:** 10.64898/2026.11.18.767696

**Authors:** Omar Molina, Leopoldo Ruiz-Huerta, Alberto Caballero-Ruiz, Ranier Gutierrez

## Abstract

Semaglutide and other GLP-1 receptor agonists are transforming obesity treatment, and patients often describe a quieting of intrusive food thoughts that motivate overconsumption of ultra-palatable foods, referred to as “food noise,” yet it remains unclear which components of feeding behavior they act upon. Here, we designed a battery of behavioral tasks that separate the perceptual, hedonic, and motivational components of glucose intake in mice. Semaglutide impaired sweet taste categorization (in a psychophysical task) and induced conditioned taste avoidance not aversion, and thus left palatability intact. Its largest effect, however, was on motivation: it selectively disrupted the active pursuit of glucose and, to a lesser extent, lipids, sparing passive intake. In a novel lick-triggered taste alternation task, two fixed sippers delivered either glucose or a second tastant (e.g., saccharin or lipids), with the tastants swapping positions after a set number of licks. This design separated effortless consumption, when the swap brought glucose directly to the mouse (passive hit), from effortful pursuit requiring crossing to the opposite sipper (active hit). A single dose of semaglutide left passive hits intact, but markedly reduced active hits: mice readily exploited glucose that came to them, yet often failed to go after it. As the drug’s effect waned, mice regained body weight by selectively overconsuming standard chow over glucose, and glucose over lipids, a shift in diet choice that persisted through the end of the experiment. Thus, semaglutide impairs the wanting, but not the liking, of glucose, providing (in mice) a behavioral correlate of the quieting of “food noise” reported by patients.

**Significance Statement:** Semaglutide preserves the hedonic liking of glucose in mice while selectively blunting its active pursuit (“wanting”), and promotes body weight rebound via a preference for chow over glucose and then lipids as the drug wanes, a profile that mirrors the suppression of “food noise” in patients. To our knowledge, our lick-triggered taste alternation task is the first behavioral assay to dissociate effortful, goal-directed caloric pursuit (“food noise” via active hits) from effortless, stationary consumption (via passive hits) and sweet taste preference in freely moving mice, offering a direct framework to dissect how anti-obesity medications reshape food seeking motivation.

## Introduction

As heterotrophs, we obtain our energy indirectly from the sun. Plants capture sunlight through photosynthesis and use that energy to synthesize glucose, which makes glucose a primary fuel for our cells and mitochondrias. In fact, we breathe to deliver oxygen to our mitochondria, and we eat to deliver the electrons stripped from glucose, whose flow to oxygen builds an H⁺ gradient (Picard and Murugan, 2025). In exchange, mitochondria convert that gradient energy into a more usable form for our body: ATP. Once ingested, however, glucose is absorbed in the gut via the sodium-glucose linked transporter 1 (SGLT1) (Sclafani et al., 2016), entering the portal vein to generate an unconscious post-ingestive signal that strongly reinforces food preference (De Araujo et al., 2020). Feeding is thus a sophisticated behavior that evolved to seek out and consume glucose-rich foods. To survive, however, omnivores must solve a harder problem: finding foods rich in glucose and other macronutrients while avoiding potential toxins (Rozin, 1976). To do so, we rely on learning and on perceptual mechanisms, among them the conscious experience of sweetness, which arises in the tongue when type II taste receptor cells (TRCs) expressing the T1R2/T1R3 heterodimer are activated (Zhao et al., 2003). Sweet taste thus serves mainly as an exteroceptive cue that predicts, but does not guarantee, the presence of glucose, because the sweet receptor is promiscuously activated by both caloric and non-caloric sweeteners. Thus, the tongue consciously predicts and the gut unconsciously confirms the presence of glucose, forming a tongue-gut-brain axis that integrates taste perception with post-ingestive glucose metabolism to shape food preference (Grove et al., 2026; Liu and Bohórquez, 2022; Tan et al., 2020). Surprisingly, how mice integrate these signals to detect, seek, and consume glucose continuously across the day-night cycle under naturalistic conditions remains largely unknown.

A second open question arises from glucose overconsumption. When intake exceeds the capacity of mitochondria to process the incoming energy flow, the surplus is stored as fat and, over the years, often leads to obesity. Glucagon-like peptide 1 (GLP-1) receptor agonists, such as semaglutide (Knudsen and Lau, 2019), are revolutionizing obesity treatment (Drucker, 2024), with patients consistently reporting a quieting of intrusive food thoughts that drive the active pursuit of ultra-palatable foods, a phenomenon termed “food noise” (Blum, 2023; Hayashi et al., 2023; Rizwan and Galyean, 2026). Yet, the precise behavioral mechanisms underlying this effect remain unclear: does semaglutide blunt sweet taste perception, induce visceral malaise and taste aversion (reducing hedonic “liking”), or selectively suppress the incentive motivation to pursue caloric rewards (“wanting”)?

Here, we introduce the Homegustometer, an automated device that records licking behavior continuously across the light-dark cycle in the home cage, and deploy a battery of behavioral tasks to dissect how a single acute dose of semaglutide and its subsequent short-term clearance reshape glucose perception, motivation, and palatability in lean, chow-fed mice. We found that semaglutide transiently impaired sweet taste categorization in a psychophysical task requiring an active perceptual report. Under naturalistic two-bottle conditions, glucose recruited post-oral mechanisms beyond sweet taste: unlike saccharin, it drove intake past the sweetness ceiling, substituted for chow calories, extended drinking into the light phase, and overrode intense sweetness (0.1% saccharin) at high (18%), though not low (3.1%), glucose concentrations. In a novel lick-triggered taste alternation task, acute semaglutide suppressed the active pursuit of glucose, while task training itself uncovered an experience-dependent enhancement in the interoceptive detection of dilute (3.1%) glucose. Crucially, a refined short-block alternation paradigm revealed that semaglutide left effortless glucose consumption (passive hits) and oromotor licking microstructure intact, while selectively suppressing the active crossings required to chase glucose (active hits). This motivational impairment generalized, albeit more weakly, to a lipid emulsion, and as the drug waned, mice regained lost body weight by preferentially overconsuming standard chow rather than glucose and lipids. Finally, semaglutide induced conditioned taste avoidance without altering hedonic palatability in a taste aversion task. Together, these findings demonstrate that semaglutide selectively impairs the “wanting,” rather than the “liking,” of specific macronutrients, providing a direct behavioral correlate for the clinical quieting of “food noise.”

## Methods

### Animals

Adult male and female mice C57BL/6 J (8-12 weeks) were used in this study. All mice were individually housed in plexiglass boxes (30 x 20 x 19 cm) in a temperature-controlled room (22 ± 2°C) with a 12-h light/night cycle (lights on at 6:00). Chip bedding, cotton nesting material, and a carton tube (enrichment) were provided to each mouse. Subjects had *ad libitum* access to standard chow diet (LabDiet 5001) composed of 4.09 kcal/g of 28.9% protein, 13.6% fat, and 57.5% carbohydrates. The Institutional Animal Care and Use Committee from Cinvestav approved experimental procedures. Unless otherwise mentioned, the equipment was cleaned daily and tastant solutions were replaced, at noon (∼12:00 h) and left available for 24 hours for *ad libitum* consumption. The Conditioned Taste Aversion (CTA) experiment was conducted during the light phase (11:00 to 15:00 h). To measure total body weight and diet consumption, mice and chow were weighed daily, at the same time of the day.

### Gustatory stimuli and drugs

Saccharin and glucose solutions were reagent-grade chemical quality (Sigma-Aldrich) and were prepared daily employing tap water. Concentrations used for glucose were 1.8, 3.1, 5.6, 10.1, and 18% w/v (100, 178, 316, 562, and 1000 mM, respectively) and for saccharin were 0.004, 0.01, 0.03, 0.1, and 0.3% w/v (0.2, 0.58, 1.71, 5, and 14.62 mM, respectively). Lipid emulsions (Fressenius Kabi) were prepared daily from Lipovenoes MCT composed of soybean oil (10 g/100 mL) and medium-chain triglycerides (10 g/100 mL) yielding 20% (1.95 kcal/mL) and diluted with distilled water to 4% (0.4 kcal/mL). Semaglutide (Ozempic, Novo Nordisk) was dissolved in 0.9% saline to achieve a final concentration of 0.067 mg/mL. Both vehicle (saline) and semaglutide (123 µg/kg) were delivered via subcutaneous injections right before initiating the dark phase (∼18:00). Lithium Chloride (Sigma-Aldrich) solution was dissolved to 200 mM in distilled water.

### The Homegustometer

The homegustometer is a custom-designed system built-in house that adapts a standard home cage for continuous monitoring of licking behavior. To dispense liquid stimuli, home cages were modified to allow the insertion of either two sippers (two bottle test and lick-triggered taste alternation task) or three sippers (water-glucose psychophysical categorization task) fixed with modelling clay. Each sipper features a drinking spout made from stainless-steel sipper tube tipped with an array of up to six 20-gauge needles, all carefully sanded at the tip and bonded together with dental acrylic cement. Each peristaltic pump (RZ1030B, Runze Fluid, China) was connected to each needle through a silicone tubing (Cole-Parmer; ID 0.062”, OD 0.125”, Wall 0.0315”). A homegustometer could operate up to 8 different pumps and thus could deliver up to 8 different tastants at a minimum volume of 1μL per lick, unless otherwise mentioned. A metal floor was attached to provide electrical ground for contact lickometers (Med Associates Inc., VT., USA).

### Water-glucose psychophysical categorization task

Six male mice were trained on a two-alternative forced-choice task to indicate whether a stimulus drop contained either glucose or water. For this behavioural paradigm, a Homegustometer configuration featuring three spouts and eight peristaltic pumps was used: six pumps delivered water and glucose cues in the central spout, while each lateral spout was connected to a single pump. Water-glucose psychophysical categorization was divided into discrimination sessions, where water and glucose 3.1% were classified, and generalization sessions, in which 1.8, 5.6, 10.1 and 18% glucose were included. The lateral spouts were separated from the central spout by a center-to-center distance of 2 cm. First, mice were trained in discrimination sessions up to 3 consecutive days reaching a performance >=70% (learning criterion); afterwards, generalization sessions were conducted until achieving learning criterion in 3 successive days.

A trial began after completing from 3 to 5 dry licks at the central stimulus spout, which triggered 2 μL cue delivery. The time elapsed from cue delivery to the last dry lick at the central spout is defined as *Center licking time*. Subsequently, mice were required to report their subjective perception of taste quality at the lateral spouts associated with dispensed cue-class glucose concentration. The lapse from the last dry lick after cue delivery at the central spout to the first dry lick in any lateral spout was termed *movement time*. Subjects were required to report a glucose cue rightward, irrespective of its intensity, or water leftward. Task contingencies were counterbalanced across subjects. If mice categorized the cue stimulus correctly, three drops of water were delivered in the corresponding lateral spout as a reward; conversely, in error trials, the water reward was omitted. The time spent licking at the lateral spouts, associated with either reward delivery or omission, was defined as *Lateral licking time*. In each experimental session, half of the trials presented water (0% cue-class glucose), while the remaining half pseudorandomly presented across the various glucose concentrations in equal proportions.

Once subjects were trained in generalization sessions, following two-days baseline period (Days 1-2), mice received the subcutaneous injection of a single semaglutide dose at the dark phase onset (∼18:00, Day 3). Behavior was monitored three additional days (Days 4-6) during the drug clearance period. A second 2-days baseline was recorded (Days 7-8) prior to vehicle (saline) administration (Day 9). Finally, a second three-day clearance phase was implemented (Days 10-12).

To filter out abnormally long trials in which the mice disengage from the task, an upper-bound outlier exclusion criterion based on a modified Tukey’s fence (*k* = 4) was applied to trial durations. Thresholds were calculated independently for each cue-class glucose (0, 1.8, 3.1, 5.6, 10.1, and 18% w/v), outcome (correct vs. incorrect), and experimental phase (discrimination and generalization), using data exclusively from sessions in which subjects reached the learning criterion (≥70% accuracy). For each stratified distribution, the cut-off threshold was defined as Q3+4×IQR, where Q3 is the third quartile and IQR is the interquartile range (Q3−Q1) (Dodge, 2008). Consequently, trial duration distributions were stratified by experimental phase, cue-class glucose, and outcome; only trials exceeding these specific limits within criterion-matching sessions were excluded from subsequent behavioral analyses.

Learning curves were estimated using the state-space modeling framework for binary observations proposed by (Smith et al., 2004). To capture performance dynamics relative to drug administration, analysis was performed over a window spanning from 150 trials prior to semaglutide/vehicle administration up to 375 trials post-administration. Additionally, to evaluate the learning trajectory throughout the mice’s stay in the Homegustometer, learning curves were calculated based on all completed trials. Finally, the learning trial was defined as the first trial in which the lower bound of the 95% confidence interval exceeds chance level and remains consistently above it for the remainder of the session (Gutierrez et al., 2010; Smith et al., 2004).

### Two-bottle preference test

All experiments were conducted in sweetness-naive cohorts of mice. Each lick was reinforced with 1µL depending on the stimulus-associated sipper. To characterize the preference for a caloric and a non-caloric sweetener relative to water, subjects were exposed in a concentration-dependent manner to either glucose (1.8, 3.1, 5.6, 10.1, and 18% w/v; 6 males and 6 females) or saccharin (0.004, 0.01, 0.03, 0.1, and 0.3% w/v; 5 males and 6 females) across six consecutive days. On day 1, water was presented in both left and right sippers to establish baseline intake. Subsequently, from days 2 to 6, daily step-wise increasing semi-logarithmic sweetener concentrations paired with water were presented to mice. Both glucose and saccharin positions were counterbalanced across days and subjects.

To evaluate the preference for a caloric over non-caloric sweetener, a first group of mice (4 males and 4 females) was presented with glucose 3.1% *vs* saccharin 0.1% w/v across four consecutive days. A second cohort (4 males and 4 females) consumed glucose 18% and saccharin 0.1% for two days, followed by two additional days of glucose 3.1% and saccharin 0.1%. Finally, a last rodent group was offered to freely lick for 0.1% saccharin and 3.1% glucose for eight days (4 males and 4 females). Solution positions were counterbalanced across days and subjects.

### The lick-triggered taste alternation task

We designed a novel behavioral paradigm to assess how animals learn to detect and pursue a caloric solution over a less caloric or less palatable alternative. For the lick-triggered taste alternation task, two block-length paradigms were evaluated, using block sizes of 50, 100, 300, 500, and 1,000 licks in the first condition, and 25, 50, 100, 200, and 400 licks in the second. For the first block-length paradigm, 50, 100, 300, 500 and 1000 licks, we used 18% glucose and 0.1% saccharin for five days and an additional session 18% glucose was replaced with 3.1% glucose (n=3 and n = 5 per sex for vehicle and semaglutide cohort, respectively). In the second block-length paradigm (25, 50, 100, 200, and 400 licks), three different variants were implemented: mice chose between 3.1% glucose and 0.1% saccharin (n=5 males and 5 females); 18% glucose and 0.1% saccharin (n=6 males and 6 females per group); 1.8% glucose and 4% lipids (n=5 males and 5 females per group). All mice were taste-naive to the stimuli, and tastants were presented in the Homegustometer. Experiments were carried out using multiple independent groups of mice, with each group containing both vehicle- and semaglutide-treated subjects (except for 3.1% glucose and 0.1% saccharin pair tastants) that received a single subcutaneous injection.

Importantly, regardless of the sapid stimuli configuration, all mice were first introduced to tastants through a prototypical two-bottle preference test implemented over a period of two days, for habituation purposes. In this phase, solutions were counterbalanced across days and subjects. From the third to the eighth day, a lick-triggered taste alternation task was implemented; for this behavioral setup, two sippers on a fixed position with an array of two needles, individually connected to a peristaltic pump through silicone tubing, delivered 1 μL of the test solution per lick. Regarding the behavioral design, unless otherwise mentioned, the solution delivery contingency was interleaved in pseudorandom blocks of 25, 50, 100, 200, and 400 licks, independently of time, over day and night. This approach allowed us to dynamically alternate which sipper delivered tastants, so that the mice had to choose the location of consumption based on what tastant was dispensed. Two properties of the design apply for every analysis below. First, the sippers themselves never move: what alternates is which sipper delivers glucose, so the animal must track the taste solution, not a place. Second, blocks advance only with licks (emitted at either sipper) and never with time: a block waits indefinitely, so time is free and the only way to end a block is to spend its lick count. For every completed block we computed a glucose preference index PI = number of licks to glucose (or preferred tastant) / block size, averaged per animal × block size × day (cells with no completed block treated as missing). A pooled PI per animal × day averaged the five block-size PIs, giving each size equal weight. For pharmacological interventions, days were grouped into three phases: baseline (day 1), drug (days 2–3, acute semaglutide or vehicle), and recovery (days 4–5). On day 5 the experiment finished 6 hours early.

### Block-transition analysis: settled origin and settled response

Each transition between consecutive completed blocks with a real swap of the glucose side defines a trial. Because the first licks of a block give the animal no opportunity to search for and exploit, both the origin and the response were defined symmetrically as the settled state: the majority side × taste of the last third of the licks of the origin and destination block, respectively. Four block states result (LeftGlu, LeftSacc, RightGlu, RightSacc), each with two possible settled responses (stay or switch), yielding eight transition types. A glucose hit is reaching the glucose-delivering state of the destination block; hits are active when the animal came from a glucose origin and had to relocate to follow it, and passive when it came from a saccharin origin and glucose arrived at the sipper where it stayed.

Additional per-trial measures were the perseverative run (licks on the previous sipper before the first lick on the opposite sipper; glucose origins only) and the eventual-hit probability (whether the animal reached glucose at any point within the destination block; never reaching it counts as censored). Metrics were computed per animal × phase (minimum 5 transitions) and matrices were pooled per group × phase. The pooled matrices were also depicted as directed graphs (edges with P ≥ 0.05, arrow width proportional to probability). Hit proportions by destination block size are pooled counts with Wilson 95% confidence intervals, compared between groups per size with Fisher’s exact test. Data are mean ± SEM unless stated otherwise.

### Real-time decomposition of block transitions

For every transition we set clock zero at the moment the contingency changes (the last lick of the origin block, which completes it and swaps the solutions) and measured cumulative times to three events: (1) the last lick on the previous sipper (detection: boundary gap plus perseverative sampling of the now-saccharin solution), (2) the first lick on the opposite sipper (arrival: crossing plus any disengagement), and (3) the start of glucose exploitation, defined as the first run of at least 5 consecutive glucose licks (falling back to the first glucose lick; transitions with no glucose lick are censored). For passive transitions only the boundary gap and the time to exploitation apply, since no relocation is needed. The same events were also expressed in lick units, displaying cumulative medians across transitions per group × phase × class; because the distributions are heavy-tailed (block completions often coincide with natural disengagements), component medians do not sum to the median total.

### Hidden Markov model

Every lick was coded as one of four categorical symbols, symbol = 2·side + taste (RS, RG, LS, LG), and paired with its log10 inter-lick interval (ILI). We fit hidden Markov models with joint categorical(4) × Gaussian(log ILI) emissions by expectation–maximization (scaled forward–backward), treating each aligned day of each animal as an independent sequence. Model order was selected by a per-animal BIC scan over K = 2–8: K = 8 was optimal in 21 of 24 animals (7 in the remaining 3), exactly doubling the K = 6 of the taste-only model by the side dimension. A single shared model (common emissions and transition matrix) was then fit to all 24 animals, states were ordered by ILI and glucose probability, and the most likely state sequence of every lick was decoded with the Viterbi algorithm. For population summaries the eight states were grouped into five classes: burst-glucose BG = {S2, S3}, burst-saccharin BS = {S1, S4}, slow-glucose SG = {S5, S6}, slow-saccharin SS = {S7}, and pause PA = {S8}. Daily state probabilities are posterior marginals (forward–backward gamma) averaged per aligned day per animal and then per group.

### Conditioned Taste Aversion (CTA)

Mice were habituated to the behavioral chambers (22 x 20 x 18 cm, Med Associates) in a sound attenuating cubicle (ENV-018-V, Med-Associates) one day before experimental procedures. CTA was conducted during the light phase (11:00 to 15:00 h) and using daily 20-minute sessions. Mice were water-deprived over 24-h preceding testing and received 1-h of supplemental water every day. Across three days, subjects were exposed to two water bottles (baseline), solutions were delivered by gravity; the next consecutive day, saccharin 0.1% w/v was presented on both bottles (acquisition); 15 minutes after session completion, rodents receive either vehicle (saline, i.p.), LiCl (150 mg/kg, i.p., from 200 mM stock) or semaglutide (123 µg/kg, s.c.). Following the 72 h-recovery period, a memory retrieval (test) was conducted, in which animals were given a two-bottle choice between water and 0.1% saccharin.

### Statistics

All data were plotted as mean ± SEM. Data analysis was performed using MATLAB 2024b (MathWorks) and GraphPad Prism 8 and JASP. Cumulative distributions were analyzed through a two-tailed Kolmogorov-Smirnov test (K-S test). For data that satisfied a normal distribution, one-, two-, and three-way repeated-measures ANOVAs, as well as two- and three-way RM mixed-effects models, were performed, followed by Tukey’s post hoc test with an alpha level of 0.05.

## Results

### Mice learn to categorize glucose versus water in a self-paced psychophysical task

To measure the subjective perceptual decision category of glucose under more ethological conditions, without the need for food deprivation, we implemented a psychophysical task in which mice sample a 2 μL drop to indicate the presence or absence of glucose at their own pace over complete 24-h sessions in a home-cage-like chamber (the Homeogustometer) equipped with one central and two lateral (left/right) spouts (**Figure 1A**). Mice initiated each trial by emitting 3–5 dry licks at the central spout, which triggered the delivery of a single cue drop of either water or glucose (50% of trials each, pseudorandomly interleaved). To obtain a reward, the animal then had to report the identity of the taste cue by licking one of the two lateral spouts; correct choices were rewarded with three drops of water, whereas incorrect choices were unrewarded. The spout–taste contingency was counterbalanced across mice (**Figure 1–figure supplement 1)**. From each trial, we computed three latencies that partition the perceptual decision into distinct epochs: consummatory cue sampling (center-licking time, from cue delivery to the last central lick), movement time (response execution, from the last central lick to the first lateral lick), and outcome evaluation (lateral-licking time, from the first to the last lick at the lateral sipper) (**Figure 1A**).

**Figure 1.**
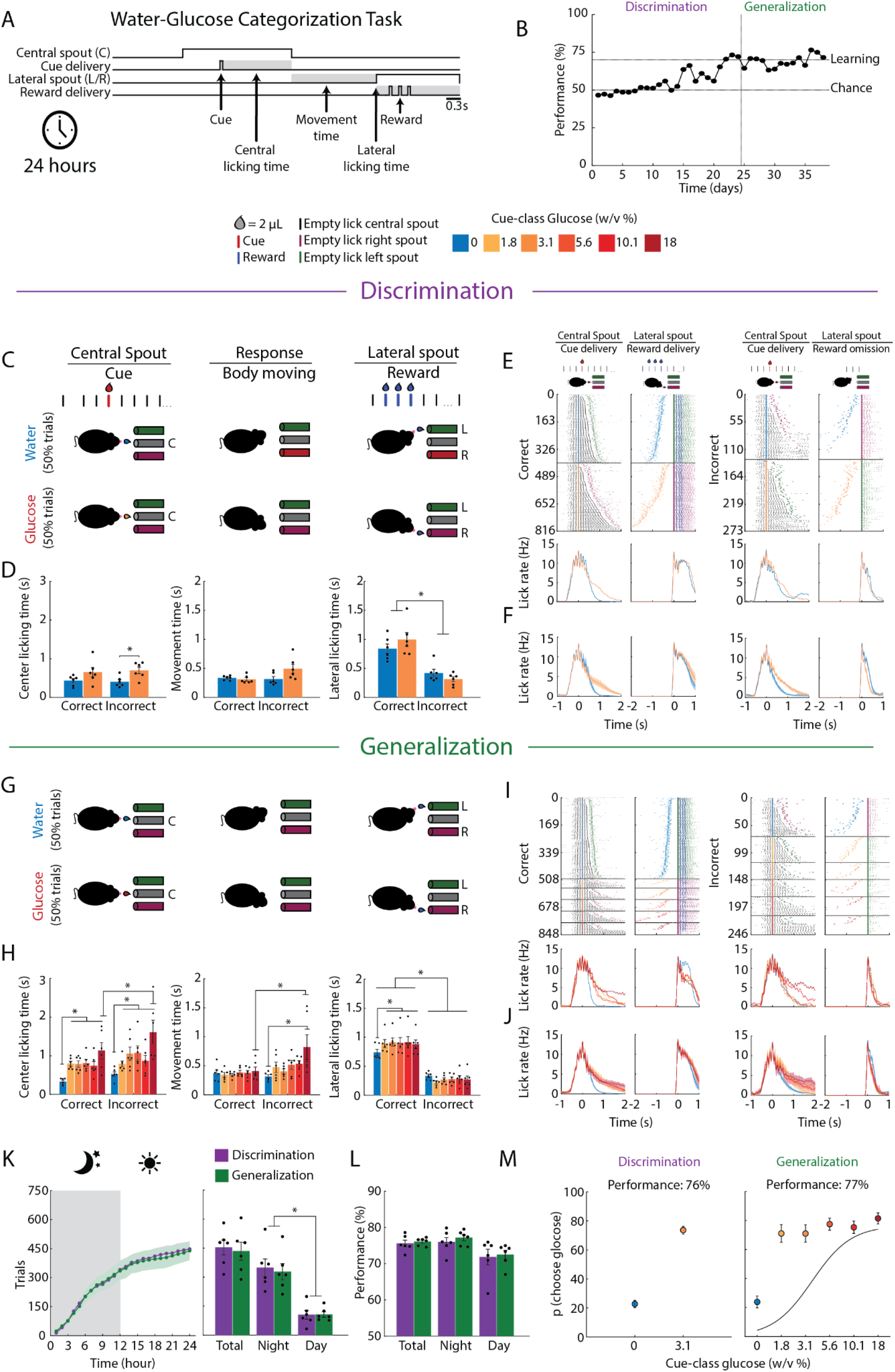
Freely moving mice learn a water–glucose psychophysical categorization task in self-paced 24-h sessions and generalize to untrained concentrations. (A). Trial structure and operational definitions. The behavioral chamber (homeogustometer) was equipped with one central and two lateral spouts, all of which were continuously available throughout the 24-h session. A trial began when the mouse performed 3–5 dry licks on the central spout, which delivered a 2 μL cue drop of either water or glucose (50% of trials each). The animal then had to choose one of the two lateral spouts; correct choices were rewarded with three drops of water, whereas incorrect trials were unrewarded. The spout–taste assignment was counterbalanced across mice (see Figure 1–figure supplement 1). Three latencies were extracted from each trial: *center licking time*, the latency to cease licking after cue delivery; *movement time*, the interval between the last lick at the central spout and the first lick at a lateral spout; and *lateral licking time*, the licking latency at the lateral spout associated with reward delivery or omission. Scale bar, 0.3 s. (B). Learning curve of a representative mouse across discrimination and generalization phases (Subject 1; in Figure 1–figure supplement 1 and Figure 2–figure supplement 1). Horizontal dashed lines indicate the learning criterion (70%) and chance level (50%); the vertical dashed line marks the transition from discrimination (purple) to generalization (green) sessions. Learning was defined as ≥70% performance on three consecutive days. Color key: cue-class glucose concentrations (0-water, 1.8, 3.1, 5.6, 10.1, and 18% w/v), coded from blue to dark red. (C). Schematic of the discrimination stage, in which mice were trained to categorize water (0%) versus a single glucose concentration (3.1% w/v). (D). Center licking time, movement time, and lateral licking time on correct and incorrect trials during discrimination sessions. (E). Licking raster plots (upper) and licking peristimulus time histograms (PSTHs, lower) from a representative mouse across three consecutive discrimination sessions, separated into correct trials (left, ending in reward delivery) and incorrect trials (right, ending in reward omission). In the rasters, ticks denote individual licks at the central (black), right (magenta), or left (green) spout; trials are sorted by trial duration and horizontal black lines delimit the cue class delivered. Within each block, the left column shows licks at the central spout aligned to cue delivery (t = 0 s; for visualization purposes only; licks in the central spout are plotted) and the right column shows licks at the lateral spouts aligned to the first lateral lick (only licks in the lateral port are shown, plus the tick for cue delivery in the central sipper). PSTH traces are color-coded by cue-class glucose concentration. (F). Population licking PSTHs (n = 6) at the central and lateral spouts for correct (left) and incorrect (right) trials, sorted by cue-class glucose concentration. (G). Schematic of the generalization phase, in which the remaining concentrations of the glucose series (1.8, 5.6, 10.1, and 18% w/v) were introduced without further training. (H). Center licking time, movement time, and lateral licking time on correct and incorrect trials during generalization sessions, plotted as a function of cue class. (I). Representative licking rasters and PSTHs during generalization; conventions as in (E). (J). Population licking PSTHs during generalization; conventions as in (F). (K). Distribution of trials across the photoperiod. *Left*, cumulative number of trials performed during the dark (moon, gray) and light (sun, white) phases in discrimination (purple) and generalization (dark green) sessions. *Right*, quantification of total, night, and day trials. Mice concentrated most trials during the dark phase. (L). Total, night, and day performance for both photoperiods. (M). Psychometric functions for the probability of choosing the glucose-associated lateral spout as a function of cue-class glucose concentration, during discrimination (left) and generalization (right). Data are plotted as mean ± SEM; filled circles represent individual animals (n = 6). Asterisks denote statistically significant differences at the 0.05 alpha level. Figure 1–figure supplement 1 shows the individual performance and licking rasters of each individual mouse. Figure 1—figure supplement 2 can be found technical details of our Homegustomer system.

Initially, during the discrimination phase, mice were trained to categorize water (0%) versus a single glucose concentration (3.1% w/v; **Figure 1C**). All mice (n = 6) acquired the task, reaching the learning criterion (≥70% correct responses on three consecutive days) after 15.5 ± 2.5 days of training (**Figure 1B**). However, acquisition speed varied considerably across subjects (range: 11–24 days), corresponding to ∼2,400–18,000 trials per mouse, performed continuously across the day–night cycle (**Figure 2—figure supplement 1**). Trial latencies carried information about the upcoming outcome: on incorrect trials, mice exhibited longer center licking times (two-way RM-ANOVA solution effect F_(1,5)_ = 6.901, *p* = 0.046) and markedly shorter lateral licking times than on correct trials (outcome effect F_(1,5)_ = 83.64, \**p* = 0.0003), the latter reflecting the rapid termination of licking upon reward omission, whereas movement time did not differ [*p* = 0.09, n.s.] (**Figure 1D**). The licking raster plots and peristimulus time histograms (PSTHs) of a representative subject (**Figure 1E**) and of the population (**Figure 1F**) confirmed this pattern: licking at the lateral spout was sustained at ∼10–12 Hz when reward was delivered but rapidly stopped within 300 milliseconds of reward omission, indicating that mice monitored reward outcome on a lick-by-lick basis, detecting reward omission as fast as 1 or 2 licks. All six mice solved the task developing different but highly repetitive licking profiles, the idiosyncrasies of these responses are shown in **Figure 1—figure supplement 1**.

We next asked whether mice had learned a categorical rule rather than simply memorizing the two anchor stimuli (0% and 3.1% glucose) used during discrimination sessions. To test this, in the generalization phase we introduced the remaining concentrations of the glucose series (1.8, 5.6, 10.1, and 18% w/v) as novel, untrained cues (**Figure 1G**). All six mice reached the learning criterion in this phase, although subjects 3 and 4 required several additional sessions to regain accuracy (**Figure 2–figure supplement 1**). Overall accuracy during generalization was indistinguishable from that during discrimination (paired t-test, t_(5)_ = 0.55, *p* = 0.6; **Figure 1M**). Moreover, the probability of choosing the glucose-associated spout increased monotonically with concentration, yielding a psychometric function consistent with mice treating glucose as a single perceptual category, irrespective of its concentration (**Figure 1M, right**). Thus, mice extrapolated the learned water–glucose boundary to concentrations never encountered in the task, a hallmark of categorical rather than exemplar-based learning. Beyond choice accuracy, the temporal structure of each trial also tracked task variables. First, center licking time increased with the glucose concentration of the cue on both correct and incorrect trials (F_(1.977, 9.887)_ = 7.325, *p* = 0.01); water trials tended to elicit fewer licks than glucose trials, consistent with the higher palatability of glucose (Mendoza et al., 2024; Perez et al., 2013). Second, on incorrect trials, movement time was longest for the highest concentrations (F_(1,5)_ = 18.64, *p* = 0.007), suggesting that slower movements were associated with a greater likelihood of error. Finally, lateral licking time dissociated correct from incorrect outcomes across all concentrations (F_(1,5)_ = 53.84, *p* = 0.0007; **Figure 1H**), indicating that mice rapidly detected reward omission; representative and population licking dynamics are shown in **Figures 1I–J**. Because sessions were self-paced, the task also provided a read-out of the circadian organization of task engagement. As expected for a nocturnal species, mice concentrated most trials in the dark phase (**Figure 1K**). Nevertheless, accuracy was equivalent between night and day and across training stages (F_(2,10)_ = 1.88, *p* = 0.202; **Figure 1L**), indicating that performance was not confined to the circadian window of peak engagement and that psychometric reports reflected stimulus control rather than circadian fluctuations in motivation.

**Figure 2.**
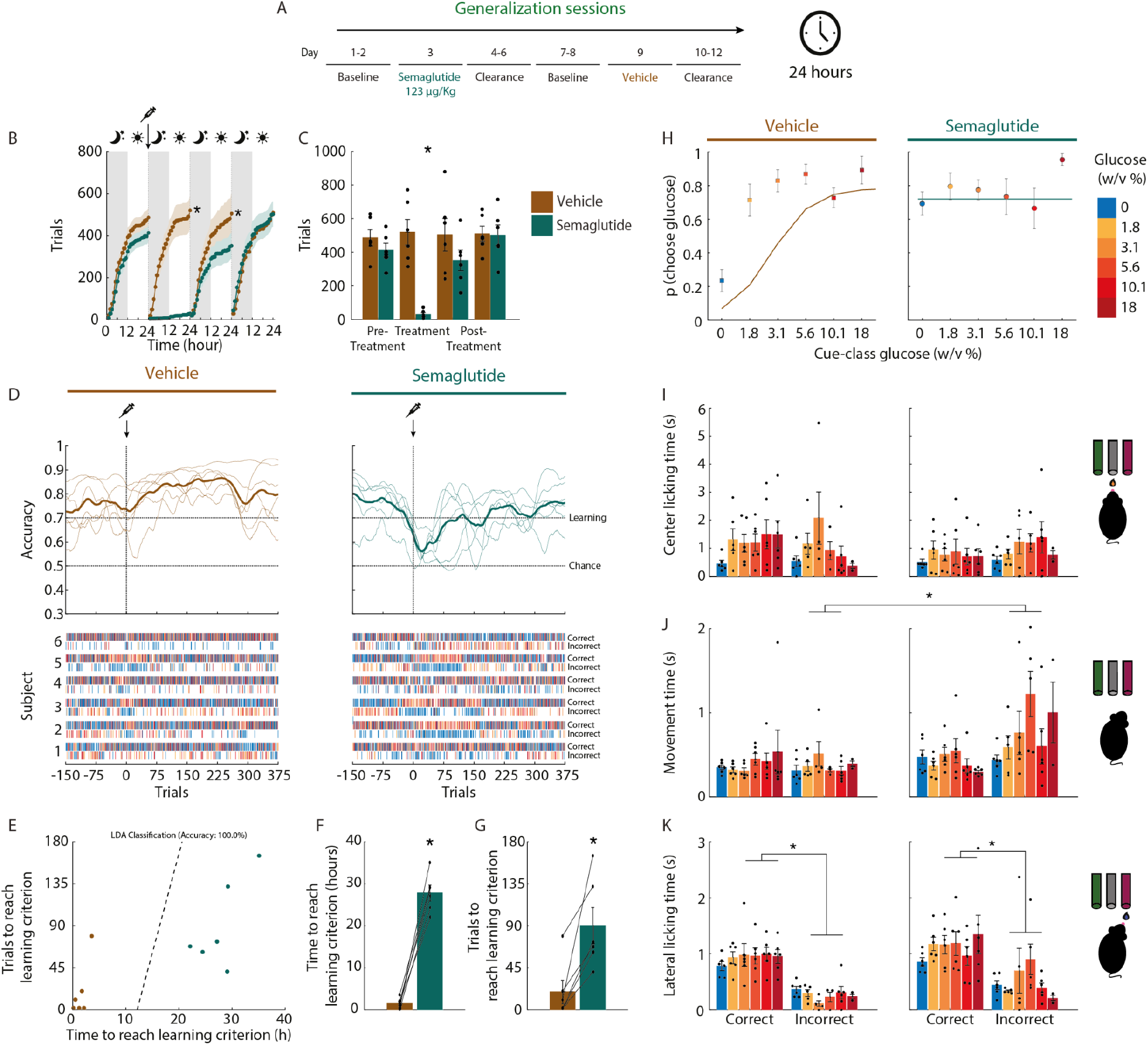
Semaglutide transiently impairs performance of the water–glucose categorization task. (A). Experimental timeline. The same six mice previously trained in the categorization task performed generalization sessions throughout and received both treatments in a within-subject design. After two baseline days (days 1–2), semaglutide (123 μg/kg, s.c.) was administered on day 3, followed by three clearance days (days 4–6). After two additional baseline days (days 7–8), the same animals received saline (vehicle) on day 9, followed by three clearance days (days 10–12). All sessions lasted 24 h and were self-paced. **(B).** Cumulative number of trials across the 24-h cycle for the baseline, treatment, and post-treatment days, for vehicle (reddish-brown) and semaglutide (dark teal). Gray and white backgrounds denote the dark (moon) and light (sun) phases; the arrow and syringe indicate treatment administration at the onset of darkness. **(C).** Total trials performed before, during, and after each treatment. Note that semaglutide, but not vehicle, reduced the number of trials on the treatment day, with subsequent recovery during clearance (post-treatment). **(D).** *Top*, task accuracy aligned to treatment administration (trial 0, arrow and syringe) for vehicle (left) and semaglutide (right). Thick traces show the group mean; thin traces show individual mice; horizontal dashed lines indicate the learning criterion (70%) and chance level (50%). Semaglutide, but not vehicle, transiently dropped accuracy toward chance, with subsequent recovery. *Bottom*, per-subject choice rasters across trials, separated into correct and incorrect trials. Colors of ticks indicate cue-class glucose concentration as in panel (**H)**. **(E).** Linear discriminant analysis (LDA) separating vehicle from semaglutide administrations based on the number of trials and the time (h) required to re-reach the learning criterion. Each dot represents one administration; the dashed line indicates the decision boundary (classification accuracy = 100%). **(F).** Time (in hours) to re-reach the learning criterion trial after each treatment. **(G).** Number of trials to re-reach the learning criterion. (**E–G).** Recovery from the semaglutide-induced impairment required substantially more time and trials than vehicle, such that an LDA could reliably classify the treatment received based on these two recovery metrics alone. **(H).** Psychometric functions for the probability of choosing the glucose-associated spout as a function of cue-class glucose concentration (0–18% w/v, color-coded) on the vehicle (left) and semaglutide (right) administration days. Note that semaglutide flattened the psychometric function around ∼0.7, indicating a loss of stimulus-guided categorization accompanied by a choice bias toward the glucose-associated spout. **(I–K).** Center licking time (**I**), movement time (**J**), and lateral licking time (**K**) on correct and incorrect trials, plotted per cue-class glucose concentration, for the vehicle (left) and semaglutide (right) days. Data are plotted as mean ± SEM; filled circles represent individual animals (n = 6, within-subject design). Asterisks denote statistically significant differences at the 0.05 alpha level. **Figure 2–figure supplement 1** shows the entire trial history of each trained mouse.

### Semaglutide transiently impairs performance of the water–glucose categorization task

Having established a self-paced psychophysical readout of sweet-intensity categorization, we asked whether a single dose of semaglutide impairs its performance. The same six trained mice performed generalization sessions continuously and received both treatments in a within-subject design: after mice achieved learning criterion (>=70%) for three consecutive days, additional stable baseline days were performed, semaglutide was administered on day 3, followed by three clearance days; after two further baseline days, the same animals received saline vehicle on day 9, followed by three clearance days (**Figure 2A**).

Semaglutide, but not vehicle, produced a near-complete suppression of trial initiation on the treatment day (treatment × drug interaction, F_(3,15)_ = 17.05, p < 0.05), with trial counts recovering to baseline during clearance (**Figure 2B–C**). Importantly, the impairment was not confined to motivation to perform: on the trials mice did complete, accuracy dropped from above criterion toward chance immediately after semaglutide administration and recovered only gradually, whereas vehicle accuracy remained unaffected (**Figure 2D**). This transient drop in performance was visible in every individual subject, both in the trial-aligned accuracy traces and choice rasters (**Figure 2D**, bottom) and in the trial-by-trial state-space learning trajectories, where the post-semaglutide trial coincided with the only sustained decrease in performance below criterion after learning (**Figure 2—figure supplement 1**).

To quantify behavioral recovery, that is, when mice regained their ability to categorize water and glucose perceptually, we measured the time and total number of trials required to re-reach the learning criterion after injection. Importantly, recovery from semaglutide was markedly delayed compared with vehicle administration, requiring substantially more time (27.8 ± 1.8 vs. 1.5 ± 0.5; paired *t*-test, t_(5)_ = 13.61, \**p* < 0.001; **Figure 2F**) and a higher number of trials (90 ± 19.4 vs. 27.8 ± 1.8; t_(5)_ = 4.116, \**p* = 0.009; **Figure 2G**). Furthermore, a linear discriminant analysis (LDA) on these two metrics alone classified the treatment received with 100% accuracy (**Figure 2E**), demonstrating that the behavioral impairment induced by semaglutide is robust and is exhibited in all individuals.

The psychometric functions revealed the nature of the impairment (**Figure 2H**). On the vehicle day, the probability of choosing the glucose-associated spout increased monotonically with cue concentration, consistent with the trained categorization (**Figure 1M, right panel**). On the semaglutide day, the psychometric function flattened: choice probability became independent of cue-class glucose concentration, remaining near 0.7 across the entire series, including water cues. The loss of the concentration-dependent slope indicates that semaglutide impaired stimulus-guided categorization rather than producing a graded shift in the category boundary. Importantly, both center and lateral licking durations remained broadly comparable between treatment conditions across concentrations (three-way RM mixed-effects model; treatment x cue-class interaction: F_(5,25)_ = 0.704, *p* = 0.24; treatment x outcome interaction: F_(1,5)_ = 0.377, *p* = 0.56; **Figure 2I–K**), indicating that the consummatory sampling of the cue and the motor execution of the choice were preserved in the trials the animals elected to perform. However, semaglutide selectively prolonged movement time during incorrect choices, as revealed by a significant treatment x Outcome interaction (F_(1,5)_ = 11.36, \**p* = 0.01). This movement delay was independent of stimulus intensity, as evidenced by a non-significant three-way interaction (cue-class glucose x treatment x Outcome: F_(5,8)_ = 1.84, *p* = 0.21), indicating that the semaglutide-induced prolongation of movement time was selectively driven by error trials, regardless of stimulus intensity. Together, these results show that a single dose of semaglutide transiently but profoundly disrupts sweet taste categorization via two main effects: one, it suppresses the willingness to engage with the task and, second, on the trials still performed, decouples choice from the orosensory evidence, with full recovery after drug clearance, taking > 24 hours. This decoupling between sensory detection and choice revealed for the first time that semaglutide transiently but robustly impairs the active report of the subjective perception of glucose from water, in mice, perhaps by prolonging movement time.

### Beyond sweetness: the caloric value of glucose drives intake that saccharin cannot match, thus “iso-sweetness” is an illusion

Having shown that acute semaglutide transiently impairs the subjective perception of glucose in the categorization task, we next asked whether it also affects the active pursuit of glucose’s caloric value. As a first step, we characterized the licking behavior of mice in a more ethological 24-h two-bottle test, in which they drank *ad libitum* and chose between water and either caloric glucose or non-caloric saccharin, presented at a different concentration each day in ascending order.

On the baseline day (day 1), when both sippers contained water, mice licked selectively during the dark phase from both spouts (**Figures 3B, 3E**). Thereafter, mice preferred both sweeteners over water in a concentration-dependent manner, with intake significantly exceeding concurrent water consumption starting at 1.8% w/v glucose (day 2) and 0.01% w/v saccharin (day 3) (**Figures 3B–C, 3E–F**; two-way RM-ANOVA solution effect: glucose,*F*_(2.72,27.29)_ = 102. 3, *p* < 0. 0001; saccharin, *F*_(2.92,26.28)_ = 41. 31, *p* < 0. 0001; K–S test, *p* < 0. 05). Higher concentrations were required to surpass baseline (day 1) total water intake, occurring from 5.6% glucose and 0.1% saccharin onward (solution effect: glucose, *F*_(2.43,24.30)_ = 57. 74, *p* < 0. 0001; saccharin, *F*_(2.69,24.29)_= 15. 02, *p* < 0. 0001).

**Figure 3.**
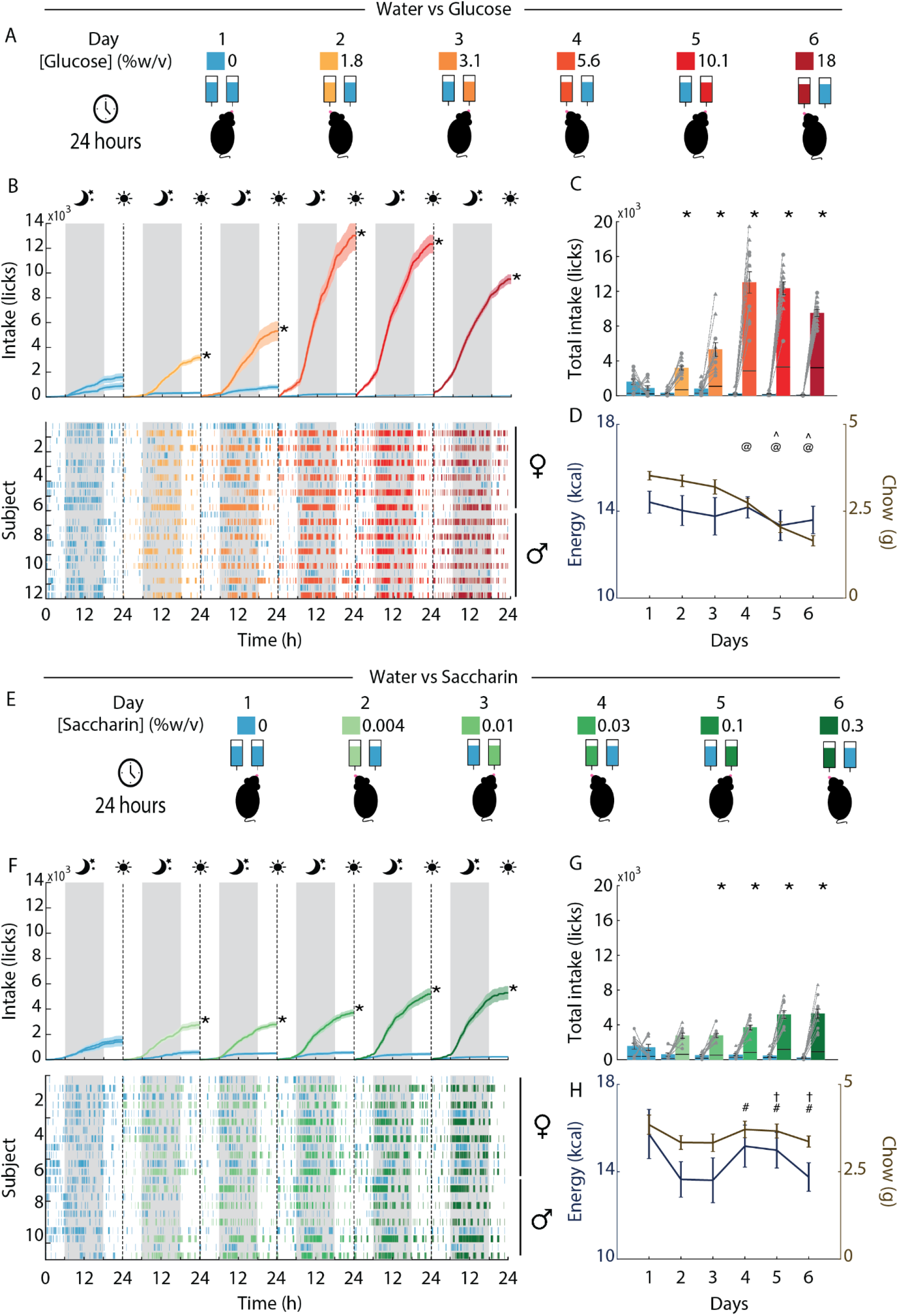
Glucose is more than sweet taste: mice prefer both glucose and saccharin over water in a concentration-dependent manner, but only glucose reaches levels beyond all saccharin concentrations, substitutes for chow calories, and extends drinking into the light phase. **(A)** and **(E).** Schematic of the two-sipper preference test. Naive mice were tested for six consecutive 24-h sessions in the Homegustometer, a home-cage system with two lickometers that continuously records licking behavior and, through peristaltic pumps, delivers a computer-controlled 1 μL drop of fluid on each lick. On day 1, both sippers delivered water (baseline). From days 2–6, mice chose between water (blue) and a sweet solution presented as an ascending semilogarithmic series, one concentration per day: glucose (1.8, 3.1, 5.6, 10.1, and 18% w/v; experiment 1) or saccharin (0.004, 0.01, 0.03, 0.1, and 0.3% w/v; experiment 2). Tastant positions were counterbalanced across days and subjects. (B). *Top:* cumulative water (blue) and glucose (color-coded by concentration as in **A**) intake across the dark (gray shading, moon) and light (white, sun) phases. Shaded bands are ± SEM. Vertical dashed lines mark the daily change in concentration and side counterbalancing. Asterisks denote significant differences in water and glucose intake over the course of a day (K-S test, *p* < 0.05). *Bottom:* lick raster of each subject (n = 12), sorted by sex (♀ top, ♂ bottom). **(C).** Total intake (licks) of water and glucose per day. Each bar is divided to show intake during the light phase (bottom segment) and dark phase (top segment). Asterisks denote significant differences in water vs. glucose within a day. Circles, males; triangles, females. **(D).** Daily energy intake from the sweet solution (kcal, left axis) and chow intake (g, right axis) in glucose-exposed mice. At and carets symbols denote significant differences in chow intake relative to day 1 for males and females, respectively; hashes and daggers denote significant differences in chow intake between the glucose group (**D**) and the saccharin group (**G**) on the same day for males and females, respectively. **(F-H).** As in **B–D**, for the saccharin experiment (n = 11). Data are mean ± SEM. Circles represent males (glucose, n = 6; saccharin, n = 5) and triangles represent females (n = 6 per group).

Despite these parallel thresholds, the two sweeteners diverged sharply in the maximum intake they sustained (**Figures 3C, 3F**). While saccharin consumption plateaued at ∼5,000 licks/day at the highest concentrations (0.1–0.3%), glucose intake continued to escalate, peaking at ∼13,000 licks/day at intermediate concentrations (5.6–10%) and remaining well above the saccharin maximum even at 18%. Thus, no concentration of saccharin ever drove intake to the levels evoked by moderate-to-high glucose concentrations.

The temporal organization of drinking also differed between sweeteners. Like water, saccharin intake remained largely confined to the dark phase across all concentrations (**Figure 3E**). In contrast, at glucose concentrations of 5.6% w/v or higher, cumulative intake curves continued to rise during the light phase, with lick rasters revealing sustained drinking invading the normally quiescent light period in both sexes (**Figures 3B–C**, light-phase bar segments). Consistent with the caloric contribution of glucose, chow intake declined progressively as glucose concentrations increased (two-way RM-ANOVA day effect: *F*_(3.14,31.38)_ = 42. 71, *p* = 0. 0001), reaching significance from day 4 (5.6%) in males (Tukey’s post hoc, *p* = 0. 0133) and day 5 (10.1%) in females (*p* = 0. 0001). Meanwhile, total daily energy intake remained stable (**Figure 3D**; *F*_(3.05,30.51)_ = 0. 84, *p* = 0. 48), indicating an isocaloric substitution of chow for glucose. No such caloric compensation occurred in the saccharin group, whose chow and total energy intake remained unchanged across days (**Figure 3G**; *F*_(2.91,26.22)_= 1. 98, *p* = 0. 14), resulting in significantly higher chow consumption compared to the glucose group across days 4–6 (three-way RM-ANOVA day × sweetener interaction: *F*_(5,95)_ = 15. 95, *p* = 0. 001). Together, these results demonstrate that while sweet taste alone establishes a preference over water, the post-oral specific value of glucose is required to drive consumption beyond the hedonic ceiling, extend drinking into the light phase, and displace solid food intake.

### When sweet taste and caloric value compete, preference tracks glucose caloric value but only at high concentrations

We next compared sweet taste and caloric value directly by letting mice choose between the two sweeteners presented simultaneously (**Figure 4**). When 0.1% w/v saccharin competed against mildly caloric 3.1% w/v glucose, mice preferred saccharin on all four days (K-S test, *p* < 0.05; **Figures 4A and 4B**), while chow and total energy intake remained stable across days (**Figure 4C**). Analysis of the licking microstructure showed that on day 1, this preference was expressed through both more numerous (1,651 vs. 751) and longer (2.3 vs. 1.8 s) licking bouts for saccharin; over subsequent days, mean bout duration converged to ∼1.5 s for both solutions, and the preference was maintained mainly by a greater number of saccharin bouts (e.g., 3,027 vs. 1,846 on day 4; **Figure 4D**), with similar initial lick rates for the two solutions.

**Figure 4.**
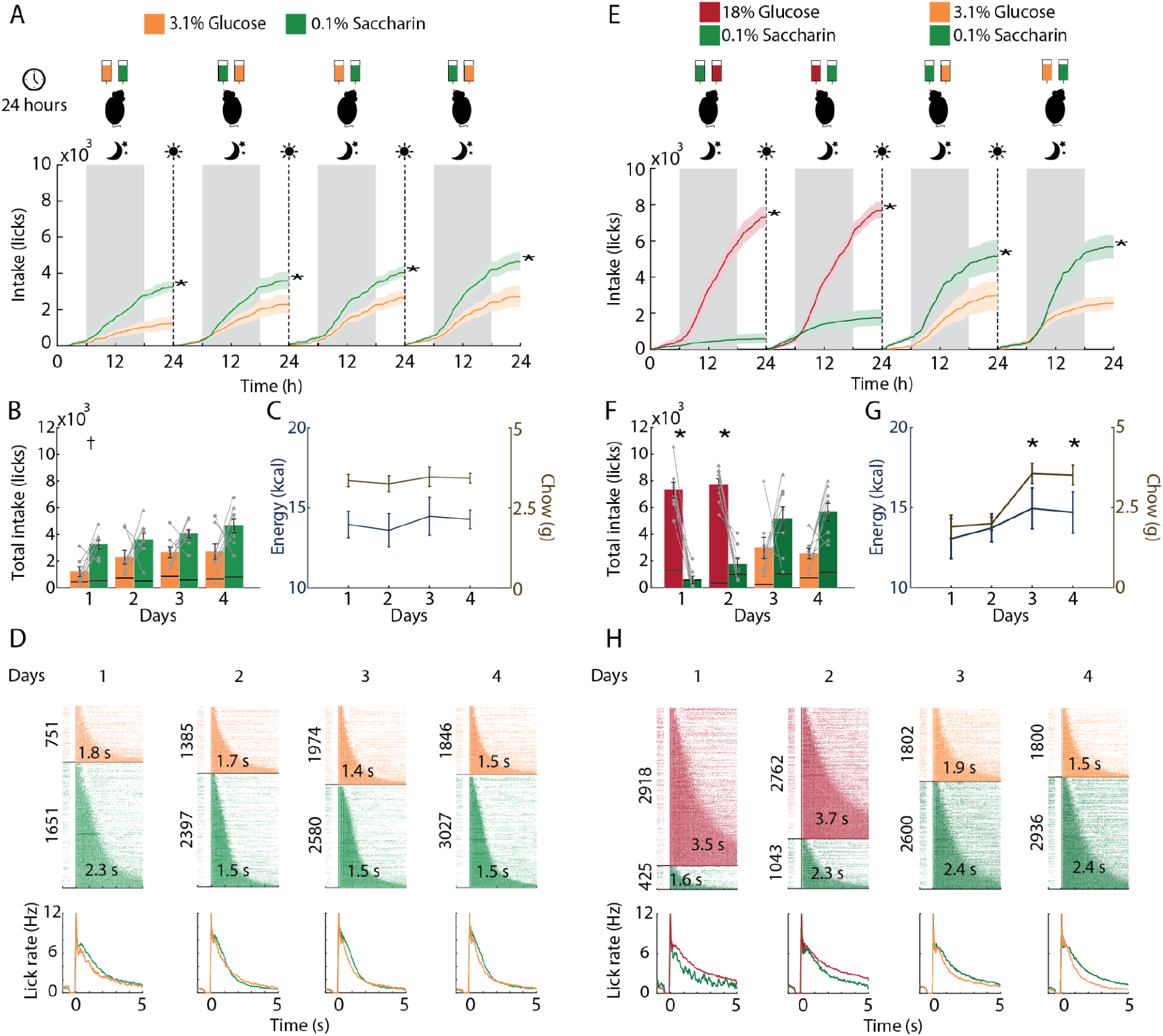
Glucose caloric value overrides sweet taste in a concentration-dependent manner: mice prefer saccharin over 3.1% glucose but reverse their choice when glucose is raised to 18%. (A). Two-sipper preference test. Mice (n = 8; 4 female and 4 male) chose between 3.1% w/v glucose (orange) and 0.1% w/v saccharin (green), the most preferred saccharin concentration in Figure 3, during four consecutive 24-h sessions in the Homegustometer. Sipper positions were counterbalanced daily (schematic). Cumulative intake (licks) is plotted across the dark (gray shading, moon) and light (white, sun) phases for each day; shaded bands are ± SEM. Vertical dashed lines mark the daily session boundary and side reversal. Asterisks denote significant differences in glucose and saccharin intake over the course of a day (K-S two-sample test, *p* < 0.05). **(B).** Total daily intake of each solution. Each bar is divided to show intake during the light phase (bottom segment) and dark phase (top segment). Circles, males; triangles, females. Asterisks as in **A**. **(C).** Daily energy intake from the solutions (kcal, left axis) and chow intake (g, right axis). **(D).** Licking microstructure. *Top:* Licking rasters in which each row is a single bout, sorted by duration (mean bout size in seconds is indicated in the white number); the number beside each raster gives the total number of bouts emitted for that solution on that day. *Bottom:* Peristimulus Time Histogram (PSTH) of the lick rate (licks/s) over the first 5 s of a bout, for each day and solution. **(E–H).** As in **A–D**, for a second cohort (n = 8; 4 females and 4 males) in which the caloric content of the glucose sipper was switched mid-test: mice chose between 18% w/v glucose (red) and 0.1% saccharin (light green) on days 1–2, and between 3.1% w/v glucose (orange) and 0.1% saccharin (dark green) on days 3–4. Note that preference tracks glucose concentration rather than sweetener identity: glucose is consumed preferentially at 18%, whereas saccharin regains preference once glucose is lowered to 3.1%. In panels (**F)** and (**G),** it can also be seen that 18% glucose extends drinking into the light phase and substitutes for chow calories, respectively. Data are mean ± SEM.

This sweetness-driven preference was overturned when the caloric content of the glucose option was raised. In a second cohort of naive mice offered 18% w/v glucose versus 0.1% w/v saccharin (days 1-2), preference reversed completely: mice consumed approximately 7,500 licks per day of glucose against fewer than 2,000 licks of saccharin (**Figures 4E** and **4F**), and glucose drinking again extended into the light phase. When the glucose sipper was then switched to 3.1% w/v (days 3-4), the preference immediately returned to saccharin, and both chow and total energy intake rose significantly (**Figures 4F** and **4G**), indicating that the calories previously obtained from concentrated glucose were recovered from chow. The licking microstructure mirrored this reversal: 18% glucose sustained markedly longer bouts (3.5-3.7 s vs. 1.6-2.3 s for saccharin) and a higher sustained lick rate within bouts, whereas after the switch saccharin regained the longer and more numerous bouts (**Figure 4H**). Hence, when sweet taste and caloric value compete directly, preference tracks the caloric content of the glucose solution rather than sweetener identity: non-caloric saccharin outcompetes weakly caloric glucose, but energy-dense glucose prevails over the otherwise preferred sweet alternative.

### Acute semaglutide produces marked, transient weight loss and hypophagia, transiently disrupts the active pursuit of caloric glucose in a lick-triggered taste alternation task, and reveals an enhanced interoceptive capacity to detect low-intensity caloric glucose after training

Based on our previous findings regarding the competition between the caloric value of glucose and the sweet taste of saccharin, we next evaluated whether semaglutide disrupts the active pursuit of glucose calories. To achieve this, we designed a novel behavioral paradigm termed the lick-triggered taste alternation task (**Figure 5A**). Briefly, this task consists of a dynamic, uncued swapping of tastants between two fixed sippers based on lick counts: each time subjects accumulated 50, 100, 300, 500, or 1,000 total licks across both sippers (drawn pseudorandomly), the solutions were automatically swapped between sippers. Through this experimental design, mice were required to continually distinguish the sweetness of saccharin from the caloric content of glucose.

**Figure 5.**
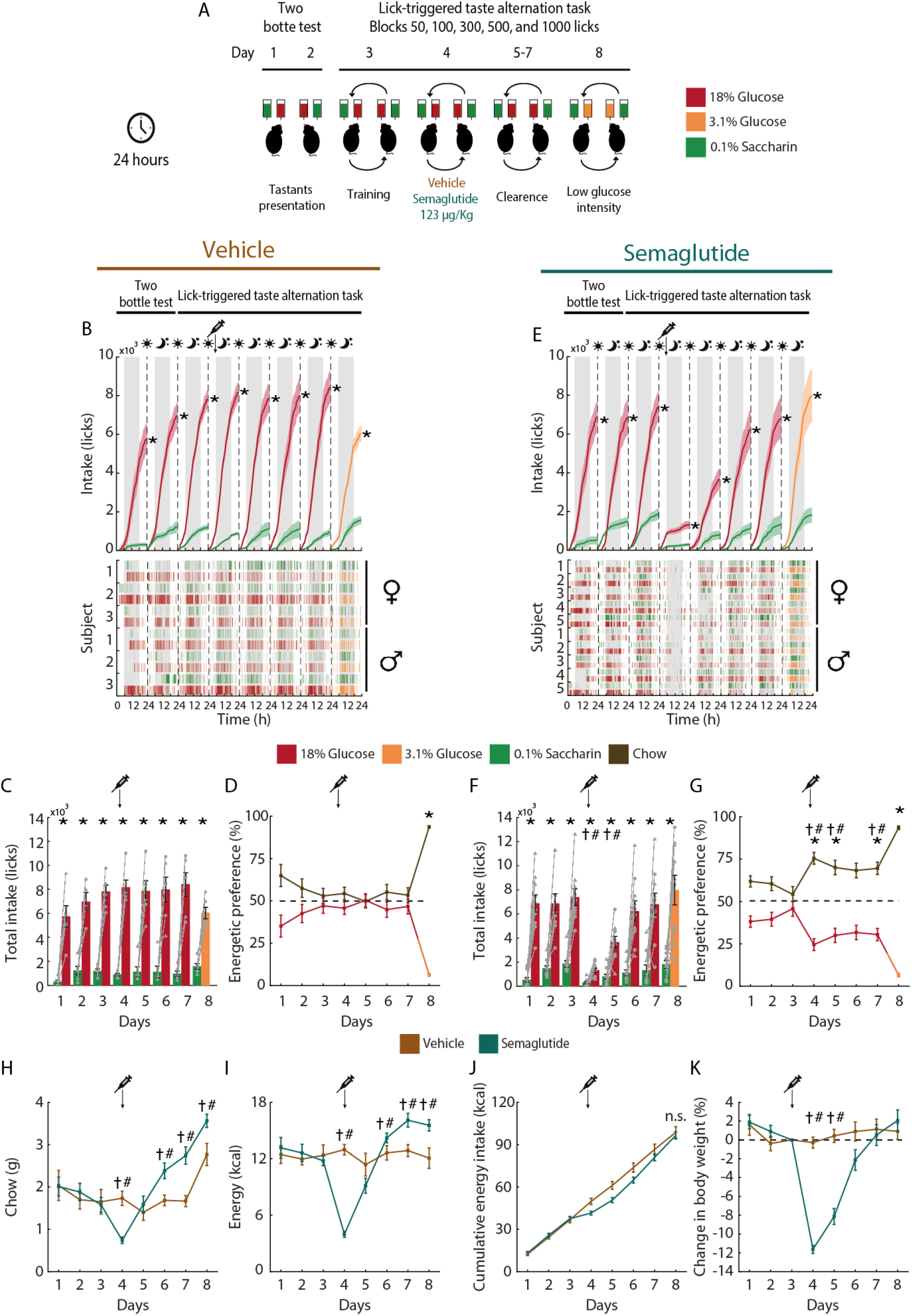
Acute semaglutide produces marked, transient weight loss and hypophagia, transiently impairs the active pursuit of caloric glucose in a lick-triggered taste alternation task, and reveals an enhanced interoceptive capacity to detect low-intensity caloric glucose after training. **(A)**. Experimental design. Mice were tested for eight consecutive 24-h sessions with two stationary sippers. On days 1 and 2, they received a standard two-sipper test with 18% w/v glucose (red) and 0.1% w/v saccharin (green); the solution assigned to each sipper was counterbalanced across days and subjects. On days 3 to 8, mice performed a lick-triggered taste alternation task. Both sippers stayed in fixed positions, and either sipper could deliver either solution: computer-controlled peristaltic pumps assigned glucose to one sipper and saccharin to the other and swapped this assignment after every block of 50, 100, 300, 500, or 1000 licks, drawn pseudorandomly. Because block transitions were uncued, mice had to sample both sippers to relocate glucose after each swap. Day 3 served as training; all mice reached criterion (glucose consumption equal to that of days 1 and 2) within this single session. On day 4, at 18:00 h (dark onset), mice received a single subcutaneous injection of vehicle or semaglutide (123 µg/kg). Days 5 to 7 allowed drug clearance. On day 8, 18% glucose was replaced with 3.1% w/v glucose (orange) to test whether mice still pursued the caloric solution when its sensory intensity was low. **(B, E)**. Cumulative licks for each solution across the dark (gray, moon) and light (white, sun) phases of the eight sessions for vehicle(left)- and semaglutide(right)-treated mice; dashed lines mark session boundaries and the syringe marks the injection (day 4). Asterisks, significant difference between solutions within a session (two-sample Kolmogorov-Smirnov test, *p* < 0.05). Bottom, lick rasters of individual subjects sorted by sex (♀ top, ♂ bottom). Mice preferred 18% glucose over saccharin in every session and, on day 8, preferred 3.1% glucose over saccharin unlike naive littermates. **(C, F)**. Total daily intake (licks) of 0.1% saccharin (green), 18% glucose (red), and 3.1% glucose (orange, day 8) for vehicle (**C**) and semaglutide (**F**) cohorts. **(D, G).** Energetic preference for both vehicle (**D**) and semaglutide (**G**) groups. Dash lines indicate indifference points; that is to say, rodents distributed equally energy intake between chow and glucose solution. **(H-K)**. Depicts chow (**H**), energy (**I**), and cumulative energy intake (**J**), as well as change in body weight (**K**) relative to prior injection day for vehicle (reddish-brown) and semaglutide (dark teal) cohorts. Syringe and arrow indicate the medication day for all panels. Data are plotted as mean ± SEM. Individual animals are represented by triangles and squares for females (n = 3 for vehicle and n = 5 for semaglutide) and males (n = 3 for vehicle and n = 5 for semaglutide), respectively. Asterisks denote differences between solutions within a day. Dagger and hashes represent differences between females and male across experimental groups, respectively. **Figure 5-figure supplement 1** depicts results from mice trained in either a two-bottle test or a lick-triggered taste alternation task comparing only 0.1% saccharin versus 3.1% glucose.

Mice (n = 6 for vehicle and n = 10 for semaglutide) first underwent a standard two-sipper test with 18% glucose versus 0.1% saccharin on days 1 and 2, followed by a single training session in the alternation task on day 3. All mice reached criterion within this single session, consuming as much glucose in the alternation task as in the two-sipper test (**Figures 5B, E**), indicating that they rapidly learned to relocate the glucose solution after each uncued swap. On day 4, at the onset of the dark phase, mice received a single subcutaneous injection of vehicle or semaglutide, followed by three clearance sessions (days 5 to 7) and a final session (day 8) in which 18% glucose was replaced with 3.1% glucose.

In vehicle-treated mice, the active pursuit of glucose was stable across all sessions (**Figures 5B, C**). Cumulative intake of 18% glucose exceeded that of saccharin in every session, including the training day and the vehicle injection day (K-S two-sample test, *p* < 0.05 for all sessions), reaching 7339 ± 833 licks for glucose versus 1041 ± 309 licks for saccharin per session (two-way RM-ANOVA, F_(1,4)_ = 98.86, *p* < 0.001). The lick rasters (**Figure 5B**, bottom) show that licking clustered in the dark phase and that mice continued to track the glucose solution across the successive swaps, with only brief sampling of the saccharin sipper. The vehicle injection itself did not alter total intake (day 4 vs. day 3, *p* = 1.0 n.s.).

In contrast, a single dose of semaglutide impaired the active pursuit of glucose on the injection day (**Figures 5E, 5F**). Total licking dropped by 83.9% and 66.7% relative to vehicle for glucose and saccharin, respectively, and the suppression was most pronounced during the dark phase immediately following the injection, when the cumulative curves remained nearly flat (**Figure 5E**, top; K-S two-sample test, *p* < 0.05). The rasters reveal that semaglutide-treated mice licked sparsely and no longer displayed the sustained bouts at the glucose sipper that characterize tracking across block swaps (**Figures 5E, 5F**, bottom). Additionally, glucose intake fell from 8162 ± 607 to 1316 ± 186 licks (F_(3.6,43.3)_ = 9.48, *p* < 0.001) and saccharin consumption dropped from 874 ± 106 to 291 ± 78 licks (F_(3.3,40.1)_ = 5.39, *p* = 0.001). The effect was transient, given that by days 6 and 7 both intake and pursuit were indistinguishable from vehicle-treated mice, total intake returned to 6212 ± 870 vs. 7967 ± 1041 licks for glucose (*p* = 0.2) and 1104 ± 242 vs 1097 ± 510 licks for saccharin (*p* = 0.98) in vehicle and semaglutide groups, respectively.

Both chow intake and energetic preference clarified the source of the body-weight rebound (**Figures 5D**, **G**, **H** and **I**). On day 3, both groups consumed the same chow amount (1.6 ± 0.1 g, *p* = 0.8) and distributed total caloric intake equally between chow (54.2 ± 0.1%) and glucose (45.7 ± 0.04%), reflected on similar energetic preference. On the injection day, semaglutide reduced daily energy intake (3.9 ± 0.3 kcal vs vehicle 12.9 ± 0.5 kcal, *p* < 0.001; **Figure 5H**); surprisingly, energetic preference was inverted by GLP-1R agonist, chow was chosen as the mainly caloric source from day 4 to 7 compared to vehicle (75.5 ± 0.07% vs 54.3 ± 0.04, respectively, *p*’s < 0.01; **Figure 5G**). Furthermore, on recovery days 6 to 8 semaglutide-treated mice increased their chow intake, resulting in a daily caloric surplus (panels **H** and **I**; F_(7,84)_ = 12.95, *p* < 0.001). In spite of semaglutide’s transient disruption of ingestive behavior, cumulative energy intake did not differ across experimental groups (**Figure 5J**, K-S test, *p* = 1). Semaglutide induced a marked, transient loss of body weight (**Figure 5K**). Relative to day 3, semaglutide-treated mice lost 11.3 ± 0.4% of their body weight by day 4, compared with 0.3 ± 0.6% in vehicle-treated mice (two-way RM-ANOVA, F_(2.57,30.91)_ = 27.47 *p* < 0.001), and this difference persisted on day 5 (*p* < 0.001). Body weight then recovered rapidly and was indistinguishable from that of controls by day 6 (*p* = 0.08); thus, the entire weight-loss episode lasted approximately two days. Thus, despite experiencing weight loss due to appetite suppression effects, semaglutide-treated mice compensated for energy deficit during clearance days through increased chow intake (a healthy food option) rather than the caloric and palatable sugar solution.

### Prior exposure to 18% glucose combined with alternation training enhances the interoceptive detection of dilute glucose

Finally, on day 8, we tested whether mice could still track the caloric solution when its sensory and energetic intensity was drastically reduced, replacing 18% with 3.1% glucose. Because 3.1% glucose provides far fewer calories than 18% glucose (0.124 vs. 0.72 kcal/mL), both groups, as expected, shifted their energetic intake toward solid chow to meet daily energy demands (**Figures 5D, G, I**). Unexpectedly, however, both vehicle- and semaglutide-treated mice preferred 3.1% glucose over 0.1% saccharin (vehicle: 6,020 ± 249 vs. 1,566 ± 481 licks; semaglutide: 7,968 ± 1,230 vs. 1,808 ± 344 licks; two-sample K-S test, both *p* < 0.01), with a non-significant trend toward even higher glucose intake in the semaglutide group (Welch’s *t*-test, *t*_(11.48)_ = 1.475, one-tailed *p* = 0.084). This preference was striking because it contrasted sharply with our previous two-bottle choice experiments: naive mice consistently preferred 0.1% saccharin over 3.1% glucose (**Figure 4A–D**), and even mice previously exposed to 18% glucose in a standard two-bottle test immediately reverted to preferring saccharin when glucose was diluted to 3.1% (**Figure 4E–H**).

How did mice in the present experiment succeed in preferring 3.1% glucose over saccharin where previous cohorts failed? We reasoned that this enhanced sensitivity could stem from three possible factors: (i) prolonged exposure (8 days) to the 3.1% glucose versus 0.1% saccharin pair itself, (ii) the cognitive demands of the lick-triggered taste alternation task alone, or (iii) the synergistic combination of prior 18% glucose exposure within the alternation task. To dissociate these possibilities, we ran two control experiments presenting 3.1% glucose and 0.1% saccharin from day 1 (**Figure 5—figure supplement 1**). First, eight consecutive days of a standard two-bottle test (**Figure 5—figure supplement 1A–D**) failed to reverse the baseline preference for saccharin, demonstrating that prolonged exposure to dilute glucose alone is insufficient for mice to prioritize its weak caloric signal over intense non-caloric sweetness.

Second, we exposed mice to two days of a standard two-bottle test followed by six days of the lick-triggered taste alternation task using only 3.1% glucose and 0.1% saccharin (**Figure 5—figure supplement 1E–H**). Rather than inducing a preference for glucose, the alternation task merely abolished the preference for saccharin, with mice licking both tastants equally. Unlike a standard two-bottle test, where spout position provides a stable spatial cue, the alternation task swaps the tastant-spout contingency continuously and unpredictably. Deprived of reliable spatial cues, mice could not link the weak post-ingestive signal of 3.1% glucose to a specific spout; instead, they alternated persistently between spouts within a block (even during large 400-lick blocks), remaining in an exploratory mode rather than exploiting either solution. This indicates that without prior experience with a strong caloric reinforcer, mice cannot rely solely on subtle taste differences between 0.1% saccharin and 3.1% glucose, and that training in the alternation task alone is insufficient to drive preference for dilute glucose. Taken together, these results demonstrate that the ability to detect and pursue a weak caloric signal emerges specifically from the combined experience of prior 18% glucose reinforcement and the attentional demands of the alternation task.

Finally, a methodological limitation of this initial task design was that block lengths reached up to 1,000 licks. Because semaglutide reduced total daily licking to ∼1,600 licks, treated mice completed only 3.1 ± 1.4 swaps on day 4 (vs. 22.3 ± 0.4 in vehicle-treated controls). This low swap count limited the temporal resolution and statistical power needed to characterize within-block glucose tracking, an issue we resolved by implementing a short-block version of the task in the next experiment (**Figure 6**).

**Figure 6.**
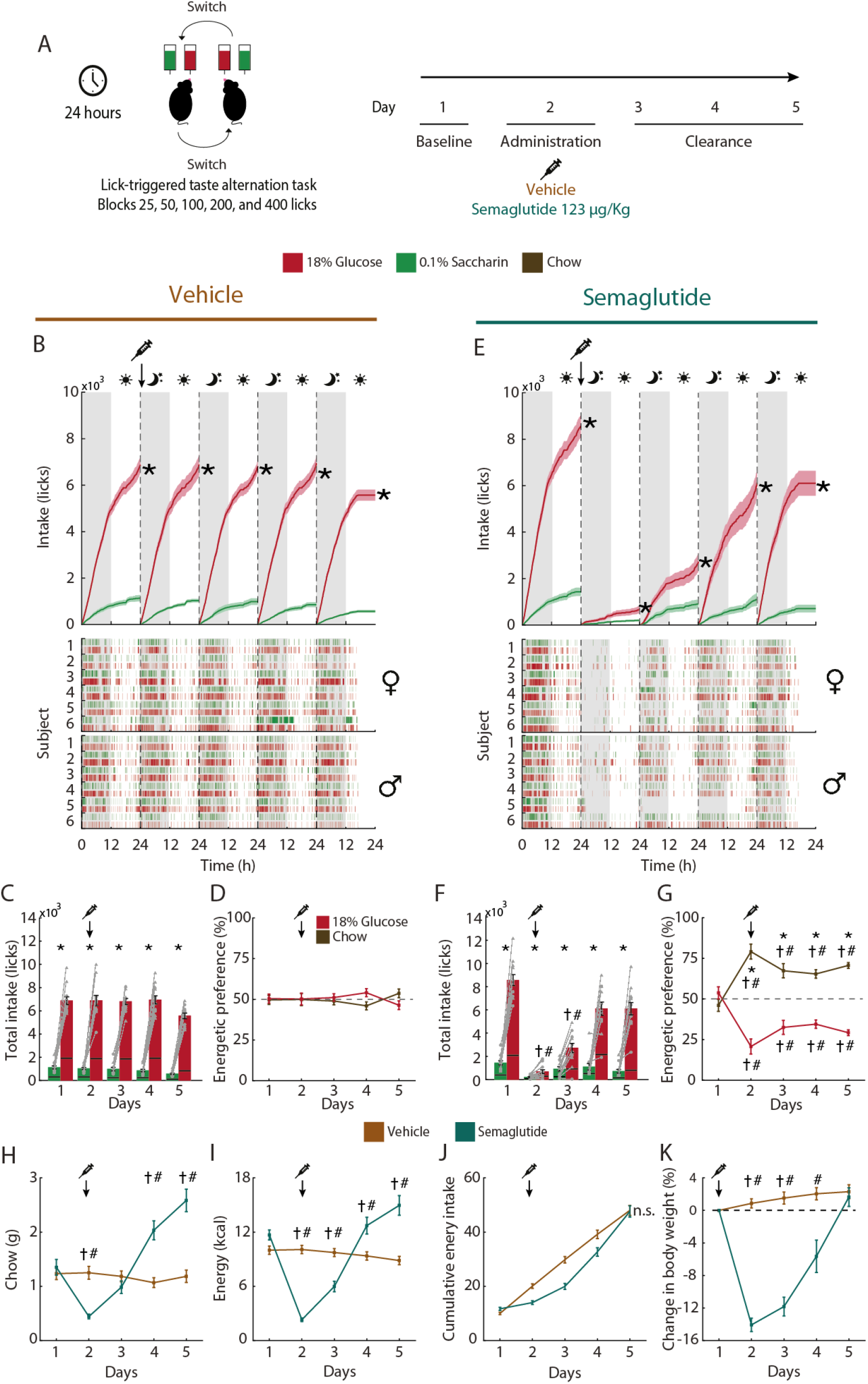
Acute semaglutide transiently impairs the active pursuit of glucose caloric value and drives a chow-directed rebound in energy intake and body weight. **(A)** Experimental design. Naive mice were tested in the lick-triggered taste alternation task with 18% w/v glucose (red) and 0.1% w/v saccharin (green), delivered through two stationary sippers in the Homegustometer, over five consecutive 24-h sessions. As in Figure 5, either sipper could deliver either solution and the assignment was swapped after a block of 25, 50, 100, 200, or 400 licks drawn pseudorandomly. The shorter blocks yielded approximately twice as many uncued swaps per session as in Figure 5, increasing the number of blocks completed under the reduced intake produced by semaglutide. Day 1 was a baseline session. On day 2, mice received a single subcutaneous injection of vehicle (brown) or semaglutide (teal) at the onset of the dark phase. Days 3 to 5 allowed drug clearance and recovery of ingestive behavior. **(B, E)** Cumulative intake (licks) of glucose and saccharin across the five sessions for vehicle- **(B)** and semaglutide-treated **(E)** mice, over the dark (gray shading, moon) and light (white, sun) phases. The syringe and arrow mark the injection; vertical dashed lines mark session boundaries; shaded bands are ± SEM. Bottom: lick raster of each subject, sorted by sex (♀ upper, ♂ lower). Note that on day 2 semaglutide-treated mice nearly stopped licking and no longer tracked glucose across swaps. On day 5 the experiment finished 6 h early. **(C, F)** Total daily intake of glucose and saccharin for vehicle **(C)** and semaglutide **(F)**. Each bar is divided into intake during the light phase (bottom segment) and dark phase (top segment). **(D, G)** Energetic preference, defined as the percentage of total daily energy intake obtained from each caloric source, the glucose solution (red) vs. chow (dark brown), with saccharin contributing no energy, for vehicle **(D)** and semaglutide **(G)**. **(H to K)** Daily chow intake **(H)**, total energy intake **(I)**, cumulative energy intake **(J),** and change in body weight as % of pre-injection body weight **(K)** for both groups. Data are mean ± SEM; n = 12 per group (6 males, circles; 6 females, triangles). Asterisks, significant differences between glucose and saccharin within a group; # and †, *p* < 0.05 between vehicle and semaglutide for the same measure in male and female mice, respectively; n.s., not significant.

### Acute semaglutide transiently impairs the active pursuit of glucose and drives a chow-directed body weight rebound

The previous experiment left a key question unresolved. Because semaglutide markedly suppressed licking and block lengths reached up to 1,000 licks, treated mice completed too few swaps to determine whether the drug had merely reduced overall intake or had specifically disrupted the active pursuit of glucose. To address this, we tested a naive cohort in a short-block variant of the task: the uncued swapping of tastants between the two fixed sippers now occurred each time subjects accumulated 25, 50, 100, 200, or 400 total licks across both sippers, drawn pseudorandomly (**Figure 6A**). This design approximately doubled the swap frequency, increasing the number of blocks completed under semaglutide while still requiring mice to continually distinguish the sweetness of saccharin from the caloric content of glucose. Mice tracked 18% w/v glucose against 0.1% w/v saccharin across five consecutive 24-h sessions, receiving a single injection of vehicle or semaglutide at the onset of the dark phase on day 2.

The cumulative intake curves provided a clear answer (**Figures 6B, E**). Both groups licked preferentially for glucose across all sessions (glucose vs. saccharin, *p* ≤ 0.0024 on all days in both groups), and in vehicle-treated mice, avid intake and tracking remained stable across days and swaps (**Figure 6B**). Thus, mice readily solved even this more demanding variant of the task. In semaglutide-treated mice, however, the injection flattened the cumulative curves for both solutions on day 2, and the lick rasters revealed that the sustained bouts at the glucose sipper that characterize tracking after each swap were replaced by sparse, scattered licks (**Figure 6E**, bottom). Thus, semaglutide did not merely reduce intake but transiently disrupted the active pursuit of glucose. Tracking reappeared partially on day 3 and was fully restored by days 4 and 5.

Total daily intake quantified the magnitude of this effect (**Figures 6C, F**). Notably, mice assigned to the semaglutide group licked more glucose than vehicle-treated mice on day 1 (8,556 ± 452 vs. 6,870 ± 321 licks, *p* = 0.005), indicating that the treated group was, if anything, more motivated by glucose at baseline; for this reason, within-group comparisons are also reported below. On day 2, semaglutide-treated mice emitted only 685 ± 152 glucose licks, a 92% decrease from their own baseline (*p* = 0.0005; vehicle: 6,885 ± 422 licks, group difference *p* < 0.0001), and 213 ± 41 saccharin licks (*p* < 0.0001 vs. vehicle). Glucose licking recovered partially on day 3 (2,718 ± 381 licks) and returned to vehicle levels by days 4 and 5 (6,108 ± 542 and 6,099 ± 513 licks, *p* ≥ 0.29). With respect to glucose licking, therefore, the suppressive effect of semaglutide was both profound and fully reversible.

Energetic preference, in contrast, did not return to baseline (**Figures 6D, G**). On day 1, both groups obtained approximately half of their daily energy from the glucose solution and half from chow (50.4 ± 2.7% in vehicle vs. 46.1 ± 4.6% in semaglutide, *p* = 0.75). Vehicle-treated mice maintained this distribution throughout the experiment (**Figure 6D**). Semaglutide-treated mice shifted their caloric intake away from glucose on day 2, as expected, but this shift did not reverse once glucose licking recovered: through day 5, their energetic preference remained displaced toward chow (**Figure 6G**). Given that glucose intake had already returned to vehicle levels, this persistent displacement necessarily reflected an increase in chow consumption.

Indeed, chow intake collapsed on the injection day and subsequently overshot baseline (**Figures 6H, I**). Groups did not differ in baseline chow intake (*p* = 0.26). On day 2, semaglutide-treated mice consumed only 0.4 ± 0.04 g of chow (vs. 1.3 ± 0.1 g in vehicle; two-way RM-ANCOVA, F_(4,79)_ = 19.52, *p* < 0.001; **Figure 6H**), and their total daily energy intake fell from 11.4 ± 0.4 to 2.3 ± 0.2 kcal (vs. 10.1 ± 0.4 kcal in vehicle; two-way RM-ANCOVA, F_(4,79)_ = 36.93, *p* < 0.001; **Figure 6I**), with only partial recovery on day 3 (6.0 ± 0.6 kcal). By days 4 and 5, however, chow intake had doubled relative to vehicle (2.0 ± 0.2 and 2.6 ± 0.2 g vs. 1.1 and 1.2 g in vehicle, respectively, *p* ≤ 0.001; **Figure 6H**), raising total energy intake significantly above control levels (12.7 ± 0.9 and 15.0 ± 1.0 kcal vs. 9.4 and 8.8 kcal in vehicle, respectively, *p* ≤ 0.001; **Figure 6I**). This compensatory hyperphagia was therefore directed specifically at solid chow rather than at the glucose solution.

This chow-directed hyperphagia was sufficient to restore cumulative energy balance and body weight (**Figures 6J, K**). The energy deficit observed under the drug was fully offset by the rebound, such that cumulative energy intake no longer differed between groups by the end of the experiment (**Figure 6J**). Similarly, groups did not differ in baseline body weight (*p* ≥ 0.98). Body weight decreased by 14.1 ± 0.8% on day 2 relative to baseline (vs. +0.9 ± 0.5% in vehicle, *p* < 0.001) and was fully restored by day 5 (**Figure 6K**). All of these effects were observed in both sexes (**Figure 6**, † females, # males). Collectively, these results indicate that a single dose of semaglutide transiently suppresses glucose pursuit and intake. As this acute effect waned, mice regained body weight not by overconsuming glucose, which merely returned to control levels, but by overconsuming chow, keeping the fraction of energy obtained from glucose below vehicle levels through day 5. Thus, beyond its acute hypophagic action, semaglutide produced a persistent shift in dietary preference away from sugar and toward balanced chow.

### Structure of the lick-triggered taste alternation task and block-by-block classification of glucose pursuit

To resolve glucose pursuit block by block, we examined the behavior of the cohort from **Figure 6** at each taste alternation (**Figure 7A**). During each 24-h session, solutions were swapped without any cue between the two fixed sippers whenever a mouse completed a pseudorandom block requirement of 25, 50, 100, 200, or 400 cumulative licks. Because the swap is uncued and sipper positions are fixed, the mouse can only detect the alternation by tasting the transition from glucose to saccharin (or vice versa) at its current sipper. We classified each block into one of three outcomes based on the mouse’s response to the new glucose location. In an active glucose hit, glucose shifted to the opposite sipper and the mouse relocated to it (typically after a few exploratory licks of saccharin), providing a direct readout of active glucose pursuit. In a passive glucose hit, the swap brought glucose to the sipper the mouse was already licking, yielding glucose intake without spatial relocation. In a miss, glucose shifted to the opposite sipper, but the mouse remained exclusively at the saccharin sipper for the entire block. By separating these outcomes, (active hits; “wanting”) specifically isolate goal-directed pursuit, which requires both taste/caloric-guided detection and spatial switching, from stationary intake (passive hits; “liking”) and failures to relocate (misses).

**Figure 7.**
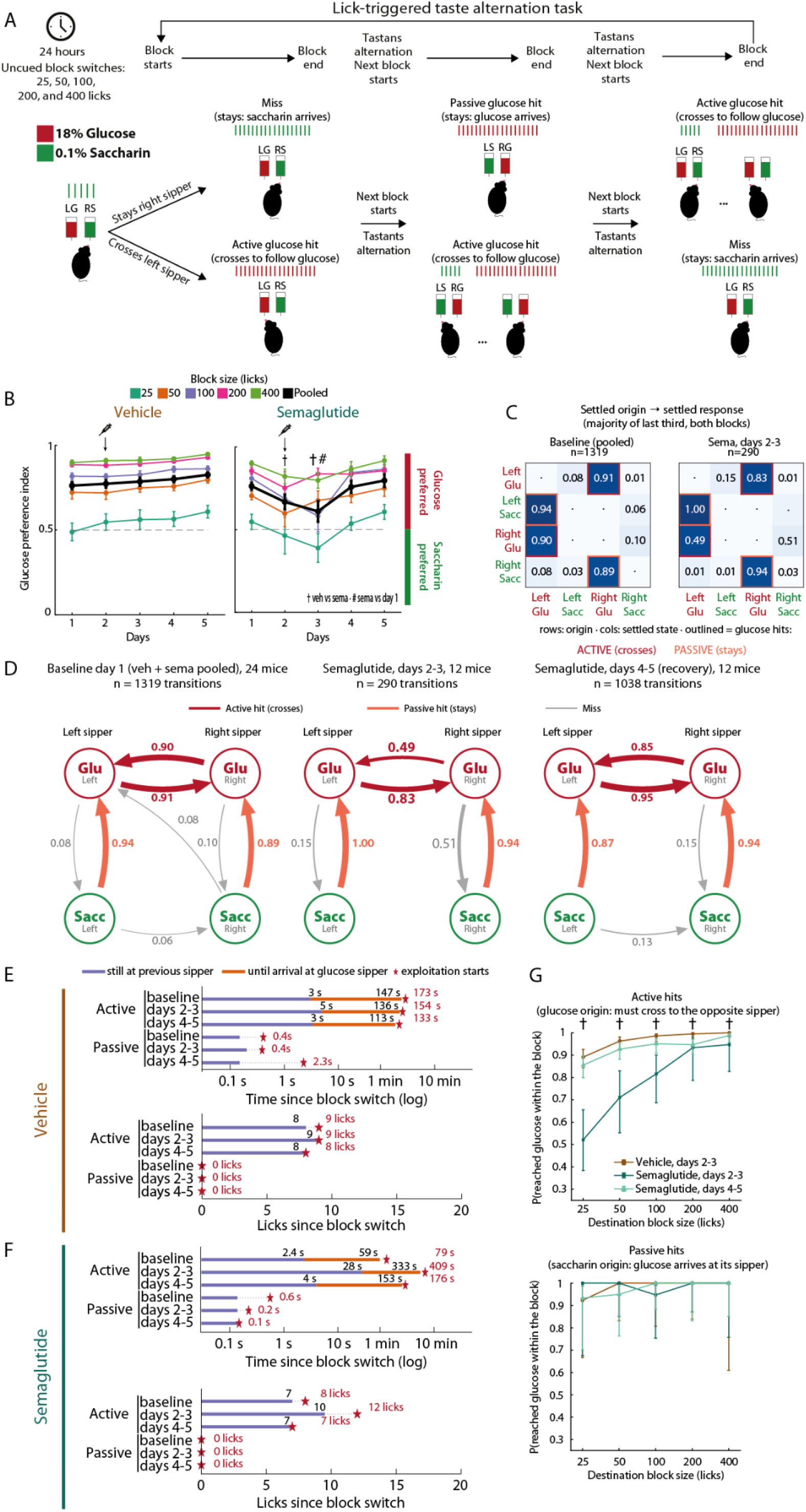
Semaglutide selectively impairs the active pursuit of glucose. Behavioral dissection of the same experiment as Figure 6 (24 mice; vehicle or semaglutide at the start of day 2; baseline = day 1, drug = days 2–3, recovery = days 4–5). **(A)** Fine grain details of structure of the lick-triggered taste alternation task and classification of block outcomes. Top: a 24-h session consists of consecutive blocks. Each block ends when the mouse accumulates a pseudorandomly drawn number of licks (25, 50, 100, 200, or 400) across both sippers; the solutions delivered by the two fixed sippers are then swapped without any cue (taste alternation) and the next block starts. Bottom: the sipper configuration in each block is denoted LG/RS (left glucose, right saccharin) or LS/RG. Red and green tick marks represent licks yielding 18% glucose and 0.1% saccharin, respectively. Each block was classified according to the behavior of the mouse relative to the new location of glucose: *active glucose hit*, the mouse crosses to the opposite sipper to follow glucose, usually after a few licks of saccharin at the sipper it was on; *passive glucose hit*, the mouse stays at the same sipper and glucose arrives (free) there as a result of the swap; *miss*, the mouse stays at the same sipper and receives saccharin, failing to relocate glucose within the block. Two example sequences of blocks are shown (upper and lower rows); ellipses indicate continuation of the block. **(B)** Per-block glucose preference index (PI, mean ± SEM) per block size (colors; 25–400 licks) and pooled across block sizes (black), across the five days for vehicle (left) and semaglutide (right). † Mann–Whitney U vehicle vs. semaglutide, # Wilcoxon vs. own day 1, *p* < 0.05; black symbols refer to the pooled PI, colored symbols to single sizes. **(C)** Settled-origin → settled-response transition matrices. Each completed block was assigned a settled state, defined as the sipper side and taste that accounted for the majority of licks in the last third of the block (LeftGlu, LeftSacc, RightGlu, RightSacc). Rows are the settled state of the origin block, columns the settled state of the next block, and each cell gives the transition probability. Left, baseline (day 1, vehicle and semaglutide groups pooled; n = 1,319 transitions); right, semaglutide days 2 and 3 (n = 290 transitions). Outlined cells are glucose hits: dark red, active (the animal crossed to follow glucose); light red, passive (glucose arrived at the sipper where the animal stayed). Dots denote transitions not observed. **(D)** The matrices in **C** displayed as directed transition graphs for baseline (day 1, both groups pooled; 24 mice, n = 1,319 transitions), semaglutide days 2 and 3 (12 mice, n = 290 transitions), and semaglutide days 4 and 5, recovery (12 mice, n = 1,038 transitions). Nodes are the four settled states, arranged by sipper (columns) and taste (rows). Arrow width is proportional to transition probability; edges with P < 0.05 are omitted. Each edge is labeled with its probability and the median time from the block switch to the first lick in the destination state. Dark red, active glucose hit; light red, passive glucose hit; gray, miss or error. The topology never changes; semaglutide re-prices the one edge that requires initiative (RightGlu → LeftGlu 0.90 → 0.49, 116 → 465 s) while the passive edges stay at ceiling in ∼0.1–0.4 s. **(E, F)** Timeline of a block transition, from the block switch to the onset of glucose exploitation (star, first run of ≥ 5 glucose licks), computed as cumulative medians across transitions, in lick units **(top)** and in real time **(bottom)**, log scale, for vehicle (**E**) and semaglutide (**F**), and for active and passive transitions in each phase (baseline, days 2 and 3, days 4 and 5). The purple segment is the time spent licking at the origin sipper after the switch and the orange segment the interval from leaving the origin sipper to the first lick at the glucose sipper. Numbers at the end of each segment give the corresponding lick count (top) or time (bottom). Statistics were computed on per-animal medians (Wilcoxon for within-group comparisons against baseline, Mann-Whitney for between-group comparisons; n = 12 per group). **(G)** Probability of reaching glucose at any point within the destination block as a function of destination block size (25 to 400 licks, log scale), for active transitions (top; glucose origin, the animal must cross) and passive transitions (bottom: saccharin origin, glucose arrives at the animal’s sipper), for vehicle days 2 and 3, semaglutide days 2 and 3, and semaglutide days 4 and 5 (recovery). Pooled proportions with Wilson 95% confidence intervals (which provide asymmetric bounds strictly constrained within 0 and 1, unlike SEM); †, Fisher’s exact test, vehicle vs. semaglutide days 2 and 3, *p* < 0.05. Because blocks advance only with licks, never with time, a failure to reach glucose means that the animal spent the block’s entire lick budget at the saccharin sipper. Number of semaglutide days 2 and 3 transitions per size: 48, 38, 49, 30, and 38 at 25, 50, 100, 200, and 400 licks.

### Glucose preference index as a function of block size dips transiently under semaglutide

The most direct indication that semaglutide affects glucose seeking is the fraction of licks in each block that yielded glucose. We therefore computed a per-block preference index (PI), defined as glucose licks divided by total licks within a block, where 0 indicates exclusive licking for saccharin, 0.5 indifference, and 1 exclusive licking for glucose (**Figure 7B**). At baseline, the PI rose monotonically with block size, ranging from approximately 0.5 in 25-lick blocks to 0.90 in 400-lick blocks. This dependence is expected from the structure of the task: after each swap, the licks spent detecting saccharin at the previous sipper and crossing to the new glucose location represent a fixed cost that consumes a large fraction of a short block, whereas it is amortized over the many glucose licks that follow in a long block. Vehicle and semaglutide groups were indistinguishable on day 1 at every block size (Mann-Whitney, all *p* ≥ 0.37; pooled: U = 68, n = 12 vs. 12, *p* = 0.84).

Semaglutide transiently lowered the PI. The pooled PI of treated mice fell from 0.77 on day 1 to 0.66 on days 2 and 3, diverging from vehicle on both days (day 2: U = 109, n = 12 vs. 12, *p* = 0.035; day 3: U = 104, n = 12 vs. 11, *p* = 0.021: one semaglutide mouse [M224] no completed blocks on day 3) and from their own baseline on day 3 (Wilcoxon W = 6, n = 11, *p* = 0.014; day 2: W = 14, *p* = 0.052; black line in **Figure 7B**). The reduction was most pronounced in the 400-lick blocks on day 3 (vs. vehicle: *U* = 109, *p* = 0.0089), which are precisely the blocks in which mice normally attain near-complete preference for glucose. This indicates that the drug degraded the efficiency with which mice pursued glucose even when the relative cost of relocating it was lowest. The PI nonetheless remained above 0.5 in most block sizes, confirming that treated mice still detected and preferred 18% glucose over 0.1% saccharin. Preference recovered completely on days 4 and 5 (pooled PI vs. vehicle, *p* ≥ 0.66; vs. their own day 1, *p* ≥ 0.15). Over the same period, vehicle-treated mice continued to improve with practice, exhibiting a higher pooled PI on days 4 and 5 than on their own day 1 (W = 5, n = 12, *p* = 0.0049 on both days), consistent with ongoing interoceptive caloric learning of the task. This simple block-level analysis thus reveals what daily intake alone could not: under semaglutide, mice could still discriminate and prefer the glucose solution, but the drug transiently reduced the fraction of their licking devoted to it, specifically dampening the active pursuit of glucose caloric value.

### Semaglutide selectively suppresses the active pursuit of glucose

The preference index quantifies how many licks within a block went to glucose, but not how mice behaved at the moment a new block began. To examine this, we characterized each completed block by the state in which the animal settled, defined as the sipper side and tastant that accounted for the majority of its licks during the final third of the block (i.e., when mice committed to exploiting a sipper: LeftGlu, LeftSacc, RightGlu, or RightSacc; **Figure 7A**), and tabulated the transitions between the settled states of consecutive blocks. Because the solutions swap at every block boundary, each transition poses one of two problems. When the origin block ends with the animal settled on glucose, the swap moves glucose to the opposite sipper: the animal must cross to keep obtaining it (active glucose hit), whereas staying yields saccharin (miss). When the origin block ends with the animal settled on saccharin, the swap brings glucose to the sipper the animal is already on: staying is sufficient (passive glucose hit), whereas crossing yields saccharin (another miss). Active hits therefore require the animal to detect the swap and act on it, whereas passive hits do not.

The resulting settled-origin to settled-response matrices (**Figure 7C**) revealed a nearly deterministic baseline policy (day 1, both groups pooled, n = 1,319 transitions): after a glucose origin, mice crossed to follow glucose (LeftGlu → RightGlu: 0.91; RightGlu → LeftGlu: 0.90), and after a saccharin origin, they stayed and received glucose (LeftSacc → LeftGlu: 0.94; RightSacc → RightGlu: 0.89). Acute semaglutide (days 2 and 3, n = 290 transitions) disrupted this policy asymmetrically. At the per-animal level, the median probability of crossing after a glucose origin collapsed from 0.92 at baseline to 0.60 on days 2 and 3, diverging significantly from vehicle controls (Mann-Whitney U = 122, n = 12 vehicle vs. 11 semaglutide, *p* = 0.0006; one semaglutide mouse [M224] was excluded for contributing fewer than five glucose-origin transitions during days 2 and 3). In the pooled matrices, this collapse was driven mainly by transitions from the right sipper (RightGlu → LeftGlu fell to 0.49, with the complementary miss RightGlu → RightSacc rising to 0.51), whereas the probability of staying after a saccharin origin remained at ceiling (0.94 to 1.00). The drug thus spared the passive acquisition of glucose and impaired only the transitions that required the animal to leave its current sipper.

The same matrices displayed as directed transition graphs make this selectivity explicit (**Figure 7D**). At baseline, virtually all traffic flowed through the four glucose-hit edges: mice either crossed to follow glucose (0.91 and 0.90) or stayed to receive it (0.94 and 0.89), with miss edges carrying at most 0.10 of the transitions. Under semaglutide, the topology of the graph did not change: the passive edges continued to operate at ceiling (1.00 and 0.94), but the active edges lost traffic, most markedly RightGlu → LeftGlu, whose probability fell by nearly half (from 0.90 to 0.49), with the lost probability mass rerouted to the perseverative miss RightGlu → RightSacc (0.51), while LeftGlu → RightGlu fell more modestly from 0.91 to 0.83.

During recovery (days 4 and 5, n = 1,038 transitions), the active edges returned to baseline levels (0.85 to 0.95) and the passive edges remained high (0.87 to 0.94). Semaglutide therefore did not alter which transitions were available, nor did it reduce glucose intake when the caloric solution arrived at the animal’s current sipper; rather, it transiently and selectively reduced the probability of the one class of transition that requires initiative: crossing to follow glucose.

### The pursuit of glucose is largely invariant in licks; semaglutide selectively stretches its duration

The transition graphs established that semaglutide reduced active transitions without affecting passive ones, but did not reveal where within the transition the deficit resided. To locate it, we aligned every transition to the block switch and reconstructed the median timeline from that moment to the onset of glucose exploitation, defined as the first run of at least five glucose licks, separately for active and passive transitions in each group and phase (**Figures 7E, F**). Each timeline was expressed in two currencies: licks emitted since the switch and elapsed time.

In lick units (**Figures 7E-F**, bottom), the twelve timelines were nearly superimposable. In active transitions, mice emitted 7 to 10 licks at the origin sipper before leaving it across every group and phase (semaglutide days 2-3 vs. baseline: W = 15.5, *p* = 0.13), and exploitation began within one or two licks of arrival at the opposite sipper (licks to exploitation onset: vehicle 9, 9, and 8 across phases; semaglutide 8, 12, and 7). The only lick-domain effect of the drug was a modest increase in licks to exploitation onset on days 2-3 (W = 7, *p* = 0.0083), which did not differ significantly between groups (U = 43.5, *p* = 0.11). In passive transitions, exploitation began at 0 licks in all conditions, including under the drug: glucose arrived in the middle of an ongoing bout and the animal simply continued licking. The sequence of licks with which mice detected the swap and relocated glucose was therefore largely preserved under semaglutide.

In real time (**Figures 7E-F**, top), however, the timelines diverged markedly. Passive transitions reached exploitation within about a second in every condition (0.1 to 2.3 s). Active transitions in vehicle-treated mice took a stable 2 to 3 min across phases (173, 154, and 133 s). In semaglutide-treated mice, the same lick sequence stretched from 79 s at baseline to 409 s on days 2 and 3, more than five times their own baseline (W = 0, *p* = 0.0005; vs. vehicle: U = 17, *p* = 0.0017). This temporal dilation affected both segments of the transition: the time spent at the origin sipper before leaving increased from 2.4 to 28 s (W = 5, *p* = 0.0049), and arrival at the glucose sipper was delayed from 59 to 333 s. By days 4 and 5, the total transition time returned toward baseline (176 s) and no longer differed from vehicle (U = 46, *p* = 0.14). Once the animal arrived at the glucose sipper, exploitation began with the very first licks in every condition. The pharmacological effect therefore resided almost entirely in the intervals between licks and between sippers, whereas the number and sequential order of the licks changed little.

### Semaglutide makes mice abandon the active pursuit of glucose when the lick budget is short

Finally, we asked how often an active transition was not completed at all by computing, for each destination block size, the probability that the animal reached glucose at any point within the block (**Figure 7G**). Interpreting a failure requires considering a key feature of the task: blocks advance only with licks emitted at either sipper, never with time. A censored transition is therefore not a timeout; rather, it means that the animal spent the entire lick budget of the block at the saccharin sipper, which completes the block, swaps the tastants back, and converts the subsequent block into a passive one. Licking out the block is thus an alternative route to glucose, and its price is the block size itself. Consequently, the pharmacological deficit scaled inversely with the price of waiting out the block (**Figure 7G**, top). Under semaglutide, the probability of reaching glucose on days 2-3 climbed from 0.52 (25/48) in destination blocks of 25 licks to 0.95 (36/38) in blocks of 400 licks, compared with 0.89 to 1.00 in vehicle controls (Fisher’s exact test: *p* = 3.4 × 10⁻⁸, 6.7 × 10⁻⁶, 1.6 × 10⁻⁵, 0.042, and 0.022 at 25, 50, 100, 200, and 400 licks, respectively), and returned to 0.86 to 0.99 on days 4 and 5. Passive hits remained at ceiling (0.92 to 1.00) across every block size and condition (all *p* = 1.00; **Figure 7G**, bottom). Notably, control mice receiving vehicle already used the strategy of staying to lick out the block (completing the block at the saccharin sipper) where it was cheap (11% of blocks of 25 licks, 24/221), and never where it was prohibitively costly (0 of 213 blocks at 400 licks). Semaglutide therefore did not impose a novel behavioral strategy; instead, it shifted the indifference point between crossing and licking out the block, such that 48%, 29%, and 18% of active transitions were censored at 25, 50, and 100 licks, respectively. The drug did not impair the animal’s knowledge of where glucose was located, nor its intake when glucose arrived passively; rather, it lowered the price at which the animal was willing to give up the active search. In other words, semaglutide decreased the incentive motivation (“wanting”) to actively pursue glucose while sparing its hedonic value (“liking”) (Berridge and Kringelbach, 2015).

### Semaglutide spares the spatial and oromotor licking responses but impairs the pause gated switch to glucose, as revealed by a lick-by-lick Hidden Markov Model

The block level analyses in **Figure 7** rest on manually defined criteria: settled states, exploitation runs, and censored transitions. We tested whether the same task structure emerges without these criteria, and whether the semaglutide induced deficit could reflect a lateralized oromotor bias. To do so, we fit a single Hidden Markov

Model (HMM) to the lick sequences of all 24 mice, describing each lick by its side and taste identity (RS, RG, LS, LG) and its log inter-lick interval (ILI). The best model (BIC, scanning *K* = 2 to 8) partitioned licking into four burst states (ILI of 125-166 ms), three slow states (203-258 ms), and one pause state of about 20 s (**Figure 7-figure supplement 1A**). Every active state was pure in symbol identity, emitting licks of a single sipper side and a single tastant. The left and right glucose states were matched in both regimes (burst, 126 vs. 125 ms; slow, 203 vs. 208 ms); the only asymmetry was the right saccharin burst state (166 ms), which had no slow counterpart, consistent with a single state absorbing both regimes at that sipper. The consummatory oromotor response is therefore essentially invariant across sides, indicating that the sipper side carries information about where glucose is located rather than reflecting a motor bias.

The transition matrix showed the same organization (**Figure 7-figure supplement 1B**). Every strong off diagonal flow linked the burst and slow states of the same symbol, and direct transitions across sides were essentially absent. Only the pause state fed both sippers, transitioning mainly into the slow glucose states (0.21 left, 0.26 right). Relocating between sippers is therefore a discrete event that passes through a pause. In emission space, the pause was the only mixed state (**Figure 7-figure supplement 1C**): its licks were split almost evenly between sides (P(right) = 0.56) but biased toward glucose (P(glucose) = 0.67). Reinitiation after disengagement is thus agnostic to side yet directed toward glucose.

Aligned to block switches, the latent classes reproduced the reversal sequence found by the manually defined analyses (**Figure 7-figure supplement 1D**): mice entered the swap bursting on saccharin at the former glucose sipper, disengaged after 10 to 15 licks (vertical dashed line), and the probability of licking the sipper that now delivered glucose rose to about 0.85 (black line). The model also exposed the block structure of the task itself, with peaks in the class probabilities at multiples of 25 licks, matching the granularity of the block sizes.

Across days, class occupancy was stable in vehicle-treated mice (**Figure 7-figure supplement 1E**, left). Semaglutide changed class occupancy in three ways (**Figure 7-figure supplement 1E**, right; per animal occupancies, Wilcoxon test vs. own day 1 and Mann Whitney test vs. vehicle, n = 12 per group). First, burst glucose occupancy fell from 0.51 to 0.27 on day 2 (W = 0, *p* = 0.0005; vs. vehicle: U = 122, *p* = 0.0043). Second, burst saccharin occupancy doubled from 0.10 to 0.23 by day 3 (W = 3, *p* = 0.0024; vs. vehicle: U = 17, *p* = 0.0017), making perseveration at the previous sipper visible as a distinct state. Third, slow glucose and pause occupancy increased on day 2 (slow-glucose, 0.31 to 0.42, W = 7, *p* = 0.0093; pause, 0.05 to 0.08, W = 5, *p* = 0.0049; pause vs. vehicle: U = 22, *p* = 0.0043). All classes returned to vehicle levels by day 5 (all *p* ≥ 0.34 vs. vehicle for the burst classes). The drug therefore changed how mice occupied each state, rather than the architecture of the model. Finally, the fraction of licks at the left sipper stayed at 0.49 to 0.50 in vehicle-treated mice across phases; under semaglutide, it shifted only transiently (0.37 on days 2-3; W = 12, *p* = 0.034 vs. own baseline) before returning to normal on days 4-5 (0.45, *p* ≥ 0.47). Together with the invariance across sides of the oromotor states, this rules out a lateralized oromotor deficit as the source of the impairment.

In sum, without any manually defined criteria, the HMM recovered a symmetric left and right architecture in which each active state belonged to a single sipper and states at opposite sippers were linked only through the pause state. Each feature has a direct behavioral meaning. First, because the model received only the raw lick sequence, this organization is intrinsic to how mice lick, not a byproduct of our analysis choices. Second, the absence of mixed active states means that mice never sample rapidly between sippers; they commit to one sipper at a time. Third, symmetry between left and right states means that mice lick with the same rhythm at both sippers, so where a mouse licks reflects a choice rather than a difference in oromotor execution. Fourth, the absence of direct transitions across sides means that changing sippers is a discrete decision: the mouse must stop licking, disengage, and reinitiate at the other sipper. Semaglutide did not alter this organization. Treated mice licked with the same rhythm at both sippers and showed no persistent side bias, consistent with a preserved representation of where glucose was delivered. What the drug changed was how mice occupied the states: greater perseveration at the previous sipper, more slow licking and pausing, and less burst licking at the new glucose sipper. In other words, semaglutide spares the structure of glucose seeking but selectively impairs the pause gated reinitiation that its active pursuit requires.

### Acute semaglutide promotes avoidance of liquid lipids, transiently impairs the active pursuit of their caloric value, and drives a chow-directed body weight rebound

Having characterized semaglutide’s effect on the preference for 18% glucose over 0.1% saccharin in the lick-triggered taste alternation task, we next asked whether the drug similarly affects the pursuit of other caloric macronutrients. We therefore tested a choice paradigm in which lipids competed directly against glucose, pairing concentrations so that lipids served as the preferred tastant: 1.8% glucose versus 4% lipids (**Figure 8A**).

**Figure 8.**
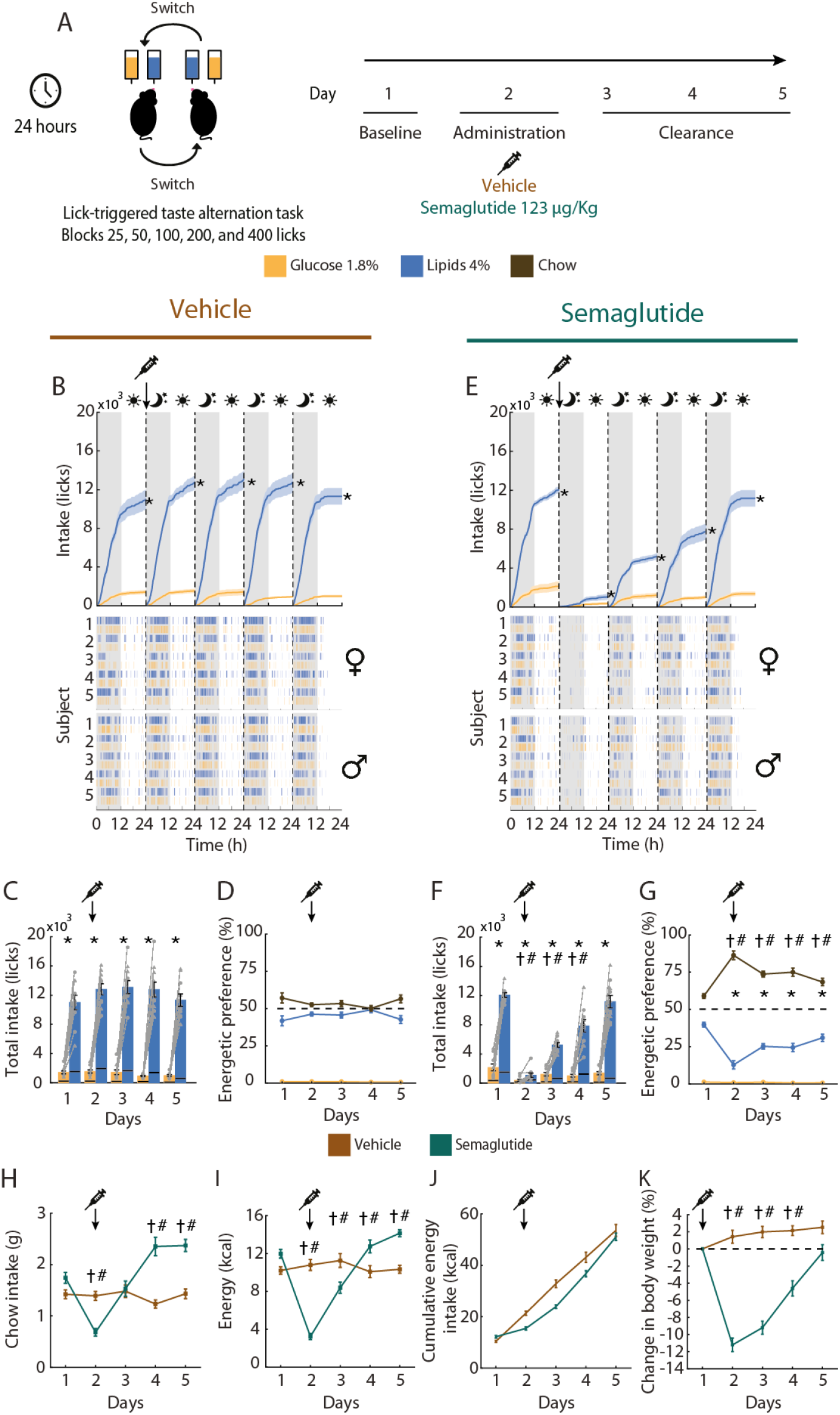
Acute semaglutide transiently suppresses lipid intake decreasing preference for a lipid solution relative to chow, followed by a compensatory rebound in chow-derived energy. (A). Experimental design. Naive mice were tested in the Homegustometer under a 24-h lick-triggered solution alternation task in which 4% lipids and 1.8% glucose (% w/v) were pseudorandomly reassigned between the two fixed sippers in blocks of 25, 50, 100, 200, or 400 licks; sipper position never changed, only the solution each one delivered. Standard chow remained available *ad libitum* throughout. Day 1, baseline; Day 2, a single subcutaneous injection of vehicle or semaglutide delivered at the onset of the dark phase (syringe and arrow); Days 3–5, clearance. Color code: orange, glucose; blue, lipids; brown, chow. **(B, E)** Cumulative licks for lipids and glucose across the five consecutive 24-h cycles in vehicle **(B)** and semaglutide **(E)** mice (top; mean ± SEM, shaded area), with the corresponding lick rasters of individual mice below, grouped by sex (♀ upper block, ♂ lower block). Gray shading indicates the dark phase, and the white background indicates the light phase; vertical dashed lines mark the day boundaries. **(C, F)** Total daily intake of lipids and glucose per day for vehicle **(C)** and semaglutide **(F)** groups. Each bar is divided to show intake during the light phase (bottom segment) and dark phase (top segment). **(D, G)** Daily energetic preference, expressed as the percentage of total caloric intake derived from each source (lipids, glucose, and chow), for vehicle **(D)** and semaglutide **(G)** groups. **(H–K)** Across-group comparison of **(H)** daily chow intake (g), **(I)** energy intake (kcal, chow plus solutions), **(J)** cumulative energy intake (kcal), and **(K)** change in body weight (%) expressed relative to the day of administration (dashed line, no change). Brown, vehicle; teal, semaglutide. Data are mean ± SEM. Circles denote males and triangles females (n = 5 per sex, n = 10 per group). \**p* < 0.05 between lipid and glucose intake within a group; # and † *p* < 0.05 between vehicle and semaglutide for the same measure for male and female mice, respectively (3-way RM-ANOVA followed by Tukey’s *post hoc*).

On day 1, baseline licking was comparable between vehicle and semaglutide groups for both tastants. On day 2, however, the GLP-1 receptor agonist drastically flattened the cumulative lick curves (**Figures 8B, E**; one-tailed two-sample K-S test, *p* is less than 0.05). Semaglutide reduced total daily intake of 4% lipids to just 8% of vehicle levels (1,087 ± 285 vs. 12,797 ± 745 licks; two-way RM-ANCOVA, *F*_(4,79)_ = 20.30, *p* is less than 0.001; **Figures 8C, F**) while concurrently suppressing 1.8% glucose licking (391 ± 132 vs. 1,538 ± 217 licks; *F*_(4,79)_ = 3.94, *p* = 0.006). Notably, the two tastants recovered at different rates (**Figure 8F**). While 1.8% glucose intake normalized by day 3 (1,223 ± 250 vs. 1,427 ± 325 licks in vehicle), 4% lipid licking recovered gradually, reaching 40% of vehicle levels on day 3 (5,237 ± 310 vs. 13,090 ± 901 licks, *p* is less than 0.001) and 61% on day 4 (7,850 ± 849 vs. 12,768 ± 1,015 licks, *p* is less than 0.001), before fully normalizing on day 5 (11,164 ± 851 vs. 11,314 ± 853 licks, *p* = 0.77). Thus, recovery from semaglutide-induced hypophagia diverged by macronutrient: intake of the preferred 4% lipid emulsion required three days to normalize, whereas 18% glucose intake in the previous paradigm recovered within two days (**Figures 5 and 6**), indicating that semaglutide induces a stronger avoidance of lipids than of glucose.

Regarding energetic distribution, vehicle-treated mice split their daily caloric intake almost evenly between solid chow (51.7 ± 1.2%) and 4% liquid lipids (47.2 ± 1.2%) across the five experimental days (**Figure 8D**; *F*_(2.4,19.6)_ = 2.79, *p* = 0.077), deriving minimal energy from 1.8% glucose (1.01 ± 0.1%, *p* is less than 0.001). In contrast, semaglutide decisively shifted energetic preference toward solid food (**Figure 8G**; *F*_(2.4,19.6)_ = 20.21, *p* < 0.001). As liquid intake collapsed on day 2, chow supplied 85.6 ± 3.1% of total calories and continued to provide 67% to 74% across days 3 to 5. Consequently, the caloric share of lipids remained below vehicle levels even on day 5 (32.5 ± 2.5% vs. 44.5 ± 2.7%), despite lipid lick counts having already normalized.

Consistent with our earlier observations (**Figures 5H–K and 6H–K**), energetic recovery was driven by solid food (**Figure 8H**). After dropping by half on day 2 (0.68 ± 0.07 vs. 1.39 ± 0.09 g in vehicle), chow intake returned to baseline by day 3 (1.53 ± 0.15 vs. 1.48 ± 0.12 g) while lipid licking remained depressed, and then significantly overshot vehicle levels on days 4 and 5 (day 4: 2.35 ± 0.18 vs. 1.23 ± 0.08 g; day 5: 2.37 ± 0.12 vs. 1.43 ± 0.09 g). This compensatory chow hyperphagia generated a daily caloric surplus on days 4 and 5 (13.0 ± 0.7 vs. 10.5 ± 0.7 kcal, and 14.5 ± 0.4 vs. 10.7 ± 0.4 kcal, respectively; **Figure 8I**) that offset the caloric deficits of days 2 and 3 (3.3 ± 0.3 and 8.5 ± 0.6 kcal), allowing cumulative energy intake to converge between groups by day 5 (51.6 ± 1.4 vs. 54.5 ± 2.4 kcal; **Figure 8J**). Mirroring chow intake, body weight dropped by 11.2 ± 0.8% within 24 h of injection (vs. +1.4 ± 0.7% in vehicle) and rebounded to within 0.4 ± 0.9% of baseline by day 5, a recovery fueled primarily by solid chow rather than the lipid emulsion (**Figure 8K**).

### Acute semaglutide only transiently and mildly slows the active pursuit of lipids

We next examined whether semaglutide altered the active pursuit of lipids to the same extent as 18% glucose (**Figure 7**). Lipid preference, defined as the fraction of licks per block directed to the lipid sipper, remained high and stable in vehicle-treated mice (0.78 to 0.86 pooled across block sizes; **Figure 9B**, left). In semaglutide-treated mice, preference dipped from 0.76 ± 0.03 on day 1 to 0.68 ± 0.04 on day 2 (**Figure 9B**, right; †*p* = 0.007 vs. vehicle, #*p* = 0.002 vs. day 1), remained slightly depressed on day 3 (0.71 ± 0.03; †*p* = 0.031), and fully normalized on days 4 and 5 (0.79 to 0.80; *p* ≥ 0.08). Although this initial drop matched the relative decrease observed for glucose preference (from 0.77 to 0.66) and was similarly driven by longer blocks reflecting exploitation rather than detection lag (decreasing from 0.84 to 0.63 in 100-lick blocks, and from 0.90 to 0.79 in 400-lick blocks), lipid preference recovered one day earlier.

**Figure 9.**
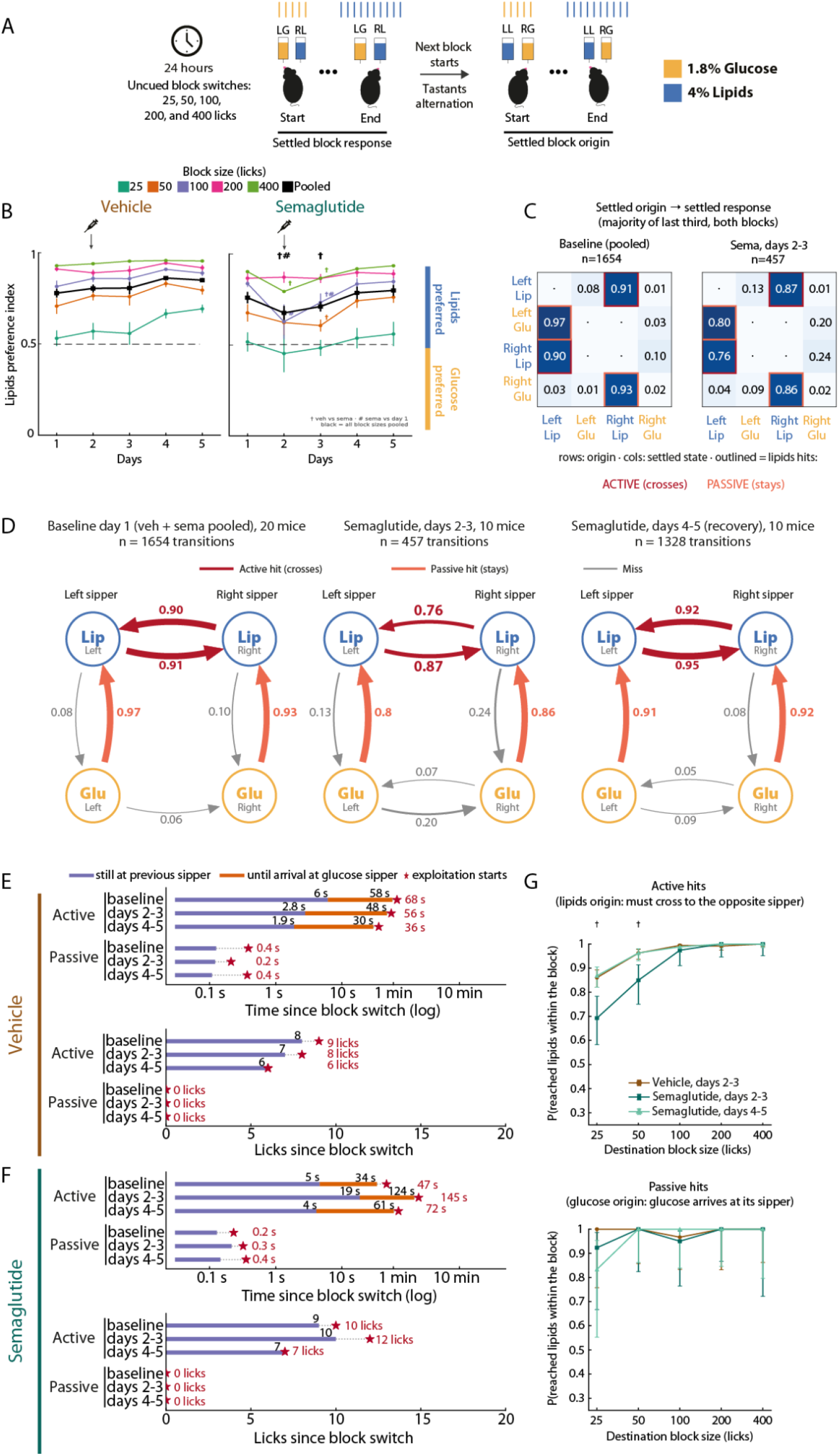
Acute semaglutide only transiently and mildly slows the active pursuit of lipids (1.8% glucose vs 4% lipids). Same mice, days, injection and color code as in Figure 8; transition definitions, timeline conventions, symbols and statistics as in Figure 7, with the lipid emulsion now the tracked solution. (**A**). Schematic representation from a block transition to another. Note that the mouse started each block consuming 1.8% glucose and, after 8-12 licks, switched to the new 4% lipids sipper. (**B**) Lipid preference (percentage of each block’s licks delivered to the lipid sipper) by block size and pooled across sizes (black) for vehicle (left) and semaglutide (right) mice; mean ± SEM, n = 10 per group. (**C)** Settled-origin to settled-response matrices at baseline (day 1, both groups pooled, 20 mice, n = 1654 transitions) and under semaglutide (days 2 and 3, 10 mice, n = 457); outlined cells, active (dark red) and passive (orange) lipid hits. (**D**) The same transitions as directed graphs, adding the recovery days (semaglutide days 4 and 5, 10 mice, n = 1328); edge labels, probability and median latency (s) from the block switch. (**E, F**) Timelines of active and passive transitions in time (upper axes, log scale) and in licks (lower axes) for vehicle (**E**) and semaglutide (**F**) mice; pooled medians. (**G**) Probability of reaching the lipids within the destination block as a function of its size, for active (top) and passive (bottom) transitions; Wilson 95% confidence intervals; † p < 0.05, Fisher’s exact test, vehicle vs semaglutide on days 2 and 3.

Settled transition matrices revealed that this transient dip shared the same structural pattern as the glucose paradigm, but with a markedly smaller amplitude (**Figure 9C**). On day 1 (both groups pooled, *n* = 1,654 transitions), mice exploiting lipids actively followed the emulsion to the opposite sipper on 0.90 to 0.91 of block swaps, matching the baseline active pursuit of glucose (0.90 to 0.91; **Figure 7D**). Conversely, mice ending a block on the glucose sipper passively stayed to exploit the newly arrived lipids on 0.93 to 0.97 of swaps. Under semaglutide (days 2-3, *n* = 457), active transition probabilities decreased moderately to 0.87 and 0.76 (with miss edges rising to 0.13 and 0.24), whereas passive transitions remained near ceiling (0.80 and 0.86). By comparison, the same pharmacological state had collapsed active transitions for 18% glucose from 0.90 to 0.49, leaving half of the transitions stranded on saccharin. Thus, the motivation to actively cross for lipids largely resisted a drug state that severely impaired the pursuit of glucose.

Transition probability graphs confirmed that the overall behavioral topology remained intact. Across all conditions, behavior was governed by the two active crossings between lipid states and the two passive self-repair edges originating from glucose (**Figure 9D**). The number of persistent licks on the depleted sipper before switching was also unaltered in both paradigms (from 7.5 to 11 licks in both experiments), confirming that semaglutide spared both the detection of contingency changes and the spatial tracking of the preferred reward.

Transition timelines further separated the behavioral components spared by the drug from those it prolonged (**Figures 9E, F**). In licks, the program was invariant, as it had been with glucose: vehicle mice crossed after 6 to 8 licks on the old sipper and began exploiting lipids at their 6th to 9th lick, semaglutide mice after 7 to 10 licks, exploiting at their 7th to 12th lick, and passive transitions required 0 licks in every condition (**Figures 9E, F**, lower axes). In time, the active transition of semaglutide-treated mice lengthened from 47 s at baseline to 145 s on days 2-3 and recovered to 72 s on days 4-5 (**Figure 9F**, upper axis), whereas vehicle transitions took 68, 56 and 36 s (**Figure 9E**, upper axis). Per animal, the time to lipid exploitation after an active transition was 162 s on days 2-3 versus 66 s in vehicle and 63 s at the same animals’ baseline (*p* = 0.006, *p* = 0.002), and the detection interval, the time still spent at the old sipper, rose from 4.8 to 17 s (*p* = 0.014, *p* = 0.002). With glucose, the same quantities had gone from 79 to 409 s and from 2.4 to 28 s. Notably, baseline active transitions were already faster for lipids than for glucose prior to treatment (68 vs. 173 s in vehicle), meaning that semaglutide imposed a smaller temporal penalty (3-fold vs. 6-fold) onto a baseline pursuit that was inherently more vigorous.

Finally, lick-budget economics explained how this modest temporal penalty translated into localized task failures (**Figure 9G**). Because blocks advance strictly by lick count, spending an entire short-block budget at the glucose sipper prevents the mouse from reaching the lipid emulsion before the next swap. On days 2 and 3, semaglutide-treated mice reduced successful active transitions into 25-lick (0.69 vs. 0.86 in vehicle) and 50-lick blocks (0.85 vs. 0.96; Fisher’s exact test, *p* = 0.0008 for both), but spared transitions into 100-, 200-, and 400-lick blocks (0.97 to 1.00 vs. 0.99 to 1.00 in vehicle; *p* ≥ 0.16), with full recovery across all block sizes on days 4 and 5 (0.87 to 1.00). In contrast, semaglutide had impaired glucose pursuit across every block length (ranging from 0.52 to 0.95 across 25- to 400-lick blocks, vs. 0.89 to 1.00 in vehicle), with several animals exhausting even the 400-lick budget on saccharin. Likewise, passive hit rates remained near ceiling in both paradigms (0.83 to 1.00 for lipids).

Together, these results show that semaglutide preserved the core lick-switching program and passive self-repair in both paradigms, while selectively inflating the latency to initiate active crossings during the first two days post-injection. The magnitude of this impairment, however, depended strongly on the target macronutrient. Despite producing acute drops in total intake and body weight comparable to those in the glucose paradigm (91% vs. 92% lick reduction; 11% vs. 14% weight loss; **Figure 8**), semaglutide reduced active crossing probabilities for lipids to only ∼0.80 (vs. 0.49 for glucose), increased transition latencies 3-fold rather than 6-fold, and restricted task failures to the shortest blocks (25 and 50 licks), where waiting out the block was cheapest. Thus, either semaglutide blunts the motivational drive for sugar more potently than for fat, or the higher caloric density of the lipid emulsion (roughly six times that of 1.8% glucose per lick) maintains its devalued incentive above the threshold required to trigger an active crossing. In other words, either semaglutide cuts sugar cravings more than fat cravings, or fat has so many calories that it is still worth the effort to go after.

### Semaglutide conditions taste avoidance (not aversion) to novel saccharin through an anorexigenic profile that spares palatability, in contrast to the canonical conditioned taste aversion produced by LiCl

The effects described so far occurred while semaglutide was pharmacologically active. Finally, we evaluated whether the interoceptive state induced by the drug also leaves a long-lasting associative trace, that is, whether a single exposure is sufficient to condition avoidance of a paired tastant, and whether such learning resembles the malaise-driven aversion produced by LiCl. Thus, to determine whether semaglutide induces visceral malaise sufficient to support associative taste learning, we used a standard conditioned taste aversion (CTA) paradigm (**Figure 10A**). Water-restricted mice received 20-min access sessions to water during baseline (days 1–3). On the acquisition day (day 4), mice were presented with a novel 0.1% saccharin solution in both bottles and, immediately after the session, received a single injection of vehicle i.p., LiCl i.p., or semaglutide s.c. (n = 8 per group). Seventy-two hours later (test, day 7), mice were given a two-bottle choice between water and 0.1% saccharin.

**Figure 10.**
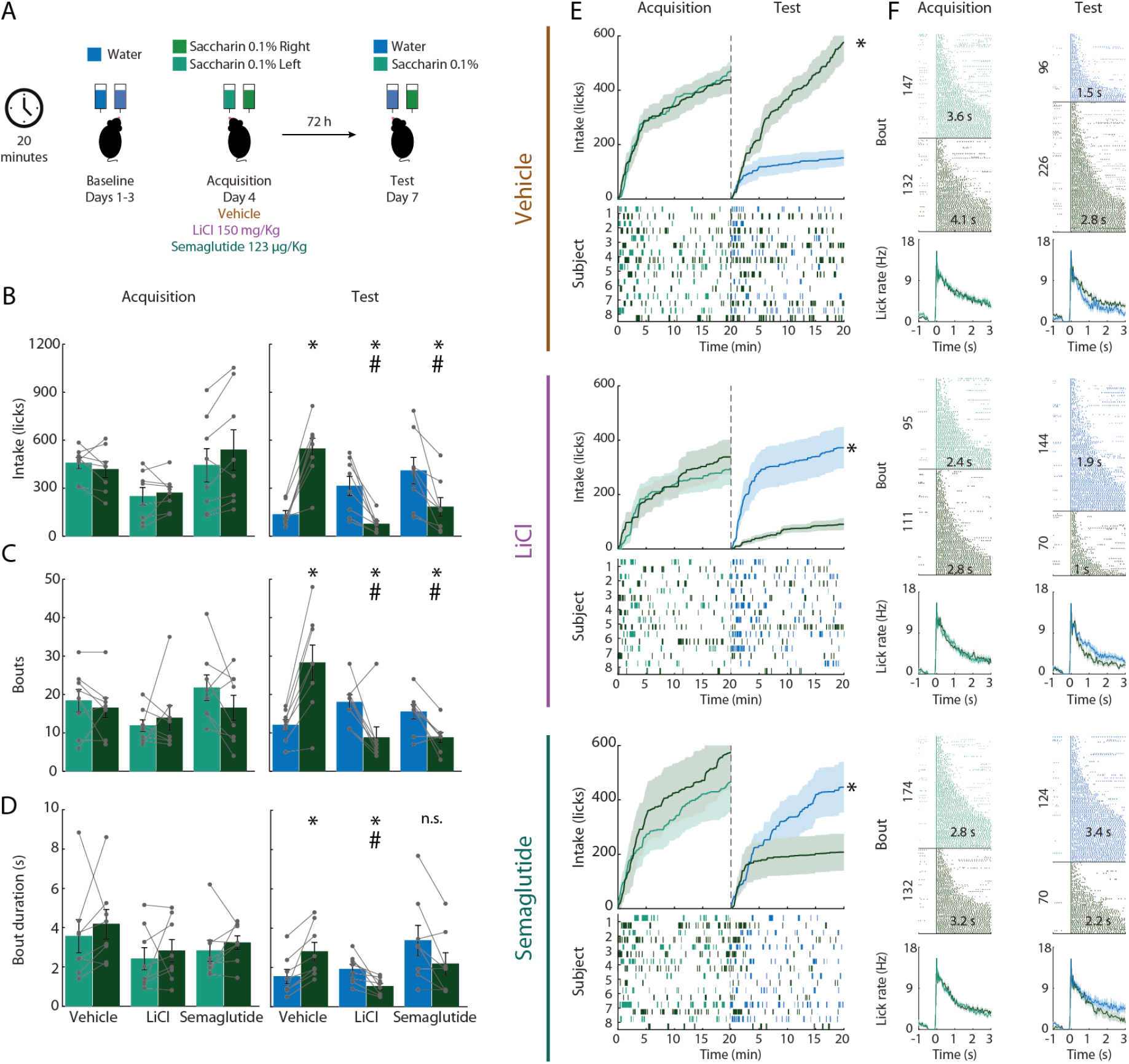
Acute semaglutide conditions taste avoidance (not taste aversion) to novel saccharin without affecting palatability. (**A**) Experimental design. Water-restricted mice were habituated to a two-bottle water setup in daily 20-min sessions during Baseline (days 1–3). On Acquisition (day 4), both bottles contained novel 0.1% saccharin (left and right positions, to control for side bias) for a single 20-min session, immediately followed by an i.p. injection of vehicle (saline), LiCl (150 mg/Kg), or semaglutide (123 μg/Kg s.c.). Seventy-two hours later (Test, day 7), mice underwent a two-bottle choice test between water and 0.1% saccharin. (**B–D**) Session-wide licking measures during Acquisition (left) and Test (right): (**B**) total intake (licks), (**C**) number of bouts, and (**D**) bout duration (seconds). Lines connect the two bottles within the same mouse. Note that saccharin intake during Acquisition did not differ among groups, indicating equivalent exposure to the conditioned stimulus prior to injection. (**E**) Cumulative lick curves (top; mean ± SEM, shaded areas) and per-subject lick rasters (bottom) across the Acquisition and Test sessions (dashed vertical line), color-coded by bottle content (green, saccharin; blue, water). Rows in (**E–F**) correspond to vehicle (top), LiCl (middle), and semaglutide (bottom). Note that at Test, vehicle-treated mice preferred saccharin over water, whereas both LiCl- and semaglutide-treated mice reversed this preference, avoiding saccharin in favor of water. (**F**) Bout-level microstructure during Acquisition (left) and Test (right). Top: raster plots of licks aligned to bout onset (each row represents one bout, sorted by bout duration; overlay is the mean bout duration per stimulus); numbers on the y-axis indicate total bouts pooled per group. Bottom: corresponding PSTHs of lick rate (Hz; mean ± SEM) aligned to bout onset. Data are mean ± SEM; symbols represent individual mice (n = 8 per group). \**p* < 0.05, saccharin vs. water within group; #*p* < 0.05, LiCl or semaglutide vs. vehicle for the same solution.

During acquisition, saccharin intake was indistinguishable across the three groups, confirming that all animals sampled the novel tastant comparably before conditioning (one-way ANOVA, treatment effect F_(2,21)_ = 2.704, *p* = 0.09, n.s.) (**Figure 10B**, left). At test, both LiCl- and semaglutide-treated mice markedly suppressed saccharin intake relative to their own acquisition levels (\**p* < 0.05) and to vehicle-treated controls (#p < 0.05), while concurrently shifting consumption toward water (**Figure 10B**, right). Vehicle-treated mice, in contrast, retained a robust preference for saccharin over water. Thus, at the level of total session intake, a single saccharin–semaglutide pairing suppressed consumption of the conditioned tastant to a degree comparable to that produced by LiCl, the gold-standard visceral malaise agent.

Analysis of licking microstructure, however, revealed that the two treatments suppressed intake through dissociable mechanisms. In both LiCl and semaglutide groups, the reduction in saccharin intake was accompanied by a significant decrease in the number of bouts directed at the saccharin spout at test (**Figure 10C**; two-way RM ANOVA, interaction F_(2,21)_ = 6.223, *p* = 0.0075), indicating diminished appetitive engagement with the conditioned tastant in both cases. Notably, bout duration dissociated the treatments: LiCl-treated mice exhibited significantly shorter saccharin bouts than vehicle-treated controls at test (**Figure 10D**; two-way RM ANOVA, days F_(1,21)_ = 29.44, *p* = 0.0001, *post-hoc* Sidak’s test #*p* = 0.0382), whereas the bout duration of semaglutide-treated mice did not differ from vehicle (*post-hoc* Sidak’s test *p* = 0.2693). Because bout duration is a well-established microstructural index of palatability, this pattern indicates that LiCl conditioning produced a genuine negative hedonic value of saccharin, the tastant became less palatable, whereas semaglutide reduced intake without degrading the palatability of the solution.

The temporal dynamics of the session reinforced this dissociation (**Figure 10E**). Vehicle-treated mice preferentially licked saccharin from the earliest minutes of the test and sustained this preference across the entire 20-min session. LiCl-treated mice displayed the mirror-image profile expected of a classical CTA: saccharin was rejected from the very first minutes of the session, with licks allocated to water from onset, consistent with the retrieval of an aversive association expressed upon initial oral contact with the tastant. In contrast, semaglutide-treated mice initially engaged with saccharin, cumulative saccharin intake rose during the opening minutes of the test, transiently paralleling the vehicle group, before consumption plateaued and preference shifted toward water as the session progressed. This delayed, within-session suppression indicates that the reduction in saccharin intake produced by semaglutide is not driven solely by an aversive, memory-based rejection of the tastant, but also by a substantial, satiation-memory-based suppression component (conditioned taste avoidance, no change in palatability, rather than pure taste aversion) that curtails ongoing intake after sampling has begun. Subject-by-subject lick rasters were consistent with this interpretation: LiCl animals showed uniform early avoidance across individuals, whereas semaglutide animals exhibited more heterogeneous profiles, with several subjects displaying appreciable early saccharin licking before disengaging.

Bout-aligned analyses recapitulated the same dissociation at the sub-second timescale (**Figure 10F**). During acquisition, within-bout lick rates (PSTHs aligned to bout onset) were comparable across groups and solutions, confirming intact oromotor execution at the time of conditioning. On test day, however, the within-bout dynamics diverged: in LiCl-treated mice, the lick rate for saccharin decayed markedly faster than for water, collapsing within the first 500 ms of bout onset, consistent with an orosensory-triggered rejection in which the taste of saccharin itself, upon contact, terminates consummatory behavior, and in agreement with the shortened bout durations of this group (**Figure 10D**). In semaglutide-treated mice, by contrast, saccharin and water lick rates were superimposable during the first second of the bout and only diverged thereafter, indicating that the initial consummatory response to saccharin remained intact and that intake was curtailed with a delay, after oral sampling had already begun.

Together, these results demonstrate that although LiCl and semaglutide produce quantitatively similar reductions in saccharin intake at test, they do so through qualitatively distinct behavioral mechanisms operating across all timescales examined. LiCl produced a canonical CTA: immediate, anticipatory avoidance of the tastant (i.e a few bouts; **Figure 10E**), shortened bouts reflecting learned hedonic devaluation (**Figure 10D**), and similar licking rates within 500 ms of tastant contact (**Figure 10F**). Semaglutide, in contrast, spared palatability, bout duration and early within-bout lick rates were indistinguishable from vehicle, and suppressed intake only after initial engagement with the tastant, both within bouts and across the session. We conclude that the intake suppression produced by semaglutide reflects a predominantly anorexigenic, satiation-like action, with at most a weak aversively associative component, whereas LiCl-induced suppression is rooted in a learned devaluation of the taste itself consisting with a frank taste aversion. This multi-timescale signature supports the interpretation that GLP-1 receptor agonism reduces feeding primarily by limiting ongoing consumption rather than by rendering foods purely aversive.

## Discussion

In this study, we designed a battery of behavioral tasks to comprehensively dissect the perceptual, palatability, and motivational components of glucose consumption, revealing how semaglutide selectively impacts the motivation to seek it. We show that acute semaglutide removes exactly the willingness to actively pursue glucose across space. Specifically, we found that semaglutide impairs the active subjective report of sweet taste categorization (in a psychophysical task), disrupts the active pursuit of glucose (and in less degree lipids; in a lick-triggered taste alternation task), and conditions taste avoidance without affecting palatability. At the level of energy balance, a single dose produced a profound but fully reversible episode, an 80% drop in daily energy intake and a 14% loss of body weight within one day, followed by a compensatory hyperphagia that restored weight within four days. Two features of that recovery are remarkable. First, the rebound was directed at chow, not at the sweet caloric solution the animals had preferred at baseline: the energetic preference for glucose remained shifted toward chow through the last day even though glucose licking itself had returned to vehicle levels. Second, throughout the episode the animals never lost the ability to sense and exploit glucose when it passively reached them, as the block-level analyses show. Clinically, patients on semaglutide do not report that food tastes worse; they report that the urge to go after it, the “food noise,” quiets down (often noting that “the freezer door is opened less frequently”). The present phenotype is the laboratory version of that report: glucose was still fully accepted when it reached the tongue; it was simply no longer worth the walk. Semaglutide left the structure of glucose seeking untouched and selectively inflated the cost of initiating its active pursuit, a pharmacological satiety that acts on the incentive salience of the pursuit of glucose, not on its content. The full recovery by days 4–5, in intake, body weight, and both the lick and time domains of the task, marks this as a reversible state imposed on an intact system: the animal keeps the map (task structure), the compass (caloric nutrient-specific tracking), and the palatability (hedonic “liking”), yet transiently loses the will to walk (motivational “wanting”).

### Advantages of the Homegustometer for naturalistic, 24-hour profiling of taste perception and ingestive behavior

Given that rodents are nocturnal, it is paramount for the neuroscience community to study their behavior in freely moving subjects under reduced human-induced stress, and more ethological day-night conditions. Thus, the Homegustometer offered three key methodological advantages by capturing taste-guided behavior continuously across the circadian cycle. First, high-resolution 24-hour monitoring revealed circadian drinking patterns that brief testing sessions inevitably miss. Beyond the expected surge in licking during the active dark phase, highly palatable and energy-dense solutions extended consumption into the inactive light phase (Petersen et al., 2023). This circadian spillover drove a substantial caloric substitution, replacing solid chow with liquid macronutrients while preserving total daily energy homeostasis (**Figures 3 to 6** and **8**). Second, the system overcame a classic bottleneck in rodent behavioral neuroscience: prolonged training and handling stress induced by human presence. Operating within a standard home cage, the Homegustometer tracked for the first time, the entire learning trajectories of a freely moving mice with minimal experimenter intervention, resolving performance both trial-by-trial during the water-glucose psychophysical categorization task (**Figure 2-figure supplement 1**) and block-by-block during the lick-triggered taste alternation task (**Figures 7 and 9**). This more ethological approach to study behavior aligns with recent advances in automated home-cage systems, which enable 24-hour auditory psychophysical testing in undisturbed mice (Yu et al., 2025). Previously used for rats (Perez et al., 2024), our 24 -hour homegustometer has been adapted into the first system to study taste psychophysics in mice. Finally, continuous monitoring provided a real-time behavioral readout of pharmacological action, allowing us to pinpoint the onset latency of semaglutide, the dynamics of intake suppression, and the macronutrient-specific trajectory of recovery during drug clearance (**Figures 2** and **5** to **9**).

### Semaglutide impacts the active report of the subjective taste perception of glucose

Although reports on GLP-1R agonist effects on gustatory active report are scarce, genetic loss-of-function studies demonstrate that GLP-1R KO mice exhibit a specific and significant reduction in lick responses during brief-access taste tests when exposed to both sucrose and sucralose (Martin et al., 2009; Shin et al., 2008; Takai et al., 2015). At the cellular level, exposure to caloric and non-caloric sweeteners, like glucose and saccharin, triggers the immediate release of GLP-1 from taste receptor cells, predominantly from T1R3-coexpressing Type II cells as well as a subset of Type III cells. In turn, GLP-1 receptors are expressed in geniculate ganglion neurons and their afferent fibers of the chorda tympani that innervate taste buds, forming a local paracrine loop in which sweeteners-induced release GLP-1 from taste buds and GLP-1 systemic administration activates sweet-sensitive gustatory nerve fibers (Takai et al., 2015), opening the possibility that GLP-1R analogues may directly modulate taste perception. Thus, GLP-1 is a hormone that recruits the tongue-gut-brain axis (Gutierrez et al., 2020).

However, in humans, the specific role of GLP-1R agonists play on taste perception remains controversial, one study using clinical psychophysical assessments through Waterless Empirical Taste Test (WETT®) demonstrated a marked reduction in taste detection and identification scores across basic taste qualities in humans under GLP-1R agonists treatment (Khan and Doty, 2025), whereas a second study found that semaglutide treatment in women with obesity and polycystic ovary syndrome alters gene expression in tongue tissue, improves overall taste active report through taste recognition scores given to a selective improvement in low intensity sweet tastants in response to sweet solution compared to distilled water, and decreases neural activation in central reward-processing regions in response to visual cues calorie-dense sweet and savory foods (Jensterle et al., 2025). Besides, it has been reported that 3 months on liraglutide treatment in type 2 diabetic obese male and female lowered sweet taste perception threshold (i.e., it enhances sweet sensitivity) during three-alternative forced-choice procedure, but reduced wanting for sweet carbohydrate foods (Brindisi et al., 2019). Data discrepancies may reflect differing sweet concentration gradients, taste assessment methods, GLP-1R agonist indications, or cohort characteristics

Our findings using the water-glucose psychophysical categorization task revealed that acute semaglutide impaired accuracy and dropped total trials performed in trained mice. In order to dissect error sources during acute semaglutide, we need to evaluate the temporal triaĺs epoch structure of our task. In the first place, these results suggest that acute semaglutide might not alter taste cues palatability given that center licking time was not affected in both correct and incorrect trials. This is in line with previous reports that GLP-1R agonists do not impact sweet taste passive oromotor responses, using a brief-access test, a task that only measures taste palatability (Acosta et al., 2026). Secondly, acute semaglutide selectively increases movement time during error trials, implying that delayed active taste reports induced by the drug may indirectly affect working memory (i.e., holding on memory the taste cue for too long), increasing decision uncertainty and errors. These data align with previous reports showing that prolonged movement times during this type of categorization task serve as a hallmark of misclassification in rats (Mendoza et al., 2024). This reinforces the notion that GLP-1R agonists by affecting incentive motivation in the response epoch, alter decision-making dynamics, though only when a volitional response is required, such as in our psychophysical task, where mice must actively report their subjective taste perception.

Finally, acute semaglutide did not alter lateral spout licking patterns; mice licked persistently during reward delivery and rapidly stopped upon omission, confirming intact sensitivity to reward outcomes. Together, normal central cue sampling paired with selectively increased movement time indicates that semaglutide spares gustatory glucose perception per se. Instead, the observed deficit in accuracy on our psychophysical task most likely reflects a targeted reduction in motivation for water rewards, promoting task disengagement under pharmacologically induced satiety.

### Lick-triggered taste alternation and semaglutide impact on motivation: The first behavioral report of “food noise” in rodents

Historically, post-oral caloric learning has been studied through four main approaches: Sclafani’s intragastric (IG) glucose infusion paradigms using saccharin-sweetened solutions (Kool-Aid) to establish post-oral “appetition” (Elizalde and Sclafani, 1990; Sclafani, 2013); de Araujo’s sucralose versus sucrose assays in sweet-blind *Trpm5* knockout mice and “dry-licking” IG paradigms showing that visceral calories alone drive striatal dopamine and foraging (De Araujo et al., 2008; Ferreira et al., 2012; Tellez et al., 2016); Using a two-bottle choice test competing glucose (or fructose) against a non-caloric sweetener mixture (0.1% sucralose + 0.1% saccharin), Sclafani and colleagues also demonstrated that glucose is the most rewarding stimulus, suggesting that mice do not merely seek calories, but rather respond to nutrient-specific, post-oral glucose sensing (Sclafani et al., 2015). Bohórquez’s discovery that gut neuropod-vagal synapses enable rapid two-bottle discrimination between sucrose and sucralose (Buchanan et al., 2022; Kaelberer et al., 2018); and Zuker’s subsequent two-bottle acesulfame-K versus glucose assays revealed a SGLT1-to-cNTS circuits specific for glucose sensing (Tan et al., 2020). Because these methods rely on invasive surgery or spatially fixed spouts, they measure cumulative consumption and preference rather than active cognitive foraging and cannot distinguish between passive and active hits within the same task. In contrast, our lick-triggered taste alternation task (saccharin versus glucose) requires mice to actively track constantly shifting tastants, unmasking interoceptive learning for dilute (3.1%) glucose. Moreover, this persistent glucose-chasing alternation parallels clinical “food noise,” the intrusive cognitive preoccupation with food (Hayashi et al., 2023). Our task revealed that semaglutide selectively quieted this active alternation, offering the first behavioral readout of “food noise” in rodents. More importantly, our results revealed that a major effect of semaglutide is on food-seeking motivation. This finding builds upon classic operant studies in rats showing that the GLP-1 receptor agonists exendin-4 (Dickson et al., 2012), reduce the breakpoint for palatable rewards in progressive ratio tasks, and directly aligns with a recent mouse study demonstrating that semaglutide both decreases the progressive ratio breakpoint and selectively delays the time required to complete each trial, using nose-poke operant task (Jones et al., 2026). Likewise, semaglutide administration also reduces progressive ratio performance in an operant wheel running task in mice (Foscue et al., 2025), suggesting it may also affect non-ingestive behaviors, such as volitional running. Taking together, our results further demonstrate that semaglutide impacts the wanting but not the liking of glucose. Our findings also suggest that this selective effect of semaglutide on incentive salience (“wanting”) partly explains its efficacy in addiction therapy by also reducing drug seeking (“cravings”) for alcohol, nicotine, and opioids (Volkow and Wargo, 2026).

### Enhanced interoceptive learning of the caloric value of dilute glucose

In our study, C57BL/6J mice initially consumed a greater volume of 0.1% saccharin than 3.1% glucose, a preference that reversed when the glucose concentration was increased to 18%. This high baseline avidity for 0.1% saccharin aligns with the presence of the sweet-sensitive Sac^b^ allele (Tas1r3) in C57BL/6J mice, which confers high T1R2/T1R3 receptor affinity and robust orosensory responsiveness to non-caloric sweeteners (Bachmanov, 2001; Fuller, 1974). Furthermore, our results parallel classic two-bottle choice studies by Valenstein, who showed that Holtzman rats prefer 0.25% saccharin over 3% glucose, but strongly favor higher glucose concentrations (6.5% and 10%) (Valenstein, 1967). Together, these observations indicate that while saccharin is initially more palatable than dilute glucose, the preference for concentrated glucose overrides pure sweet taste signaling through post-oral reinforcing effects. 3% glucose is a concentration that humans barely report as sweet (Valenstein, 1967). Crucially, we demonstrate for the first time that prior exposure to 18% glucose, combined with training in our lick-triggered alternation task, enables mice to detect and actively pursue the caloric value of dilute (3.1%) glucose. This reveals a novel form of interoceptive caloric glucose learning: requiring the animal to pay structured, active attention to gustatory cues recalibrates its interoceptive sensitivity, likely via gut–brain post-ingestive circuits, allowing it to detect the metabolic value of dilute solutions that normally fall below the hedonic threshold.

### Macronutrient-specific recovery hierarchy and chow-directed body weight rebound

A particularly notable finding of this study is the selectivity of the compensatory body weight rebound during drug clearance. Across all tasks, mice regained lost body weight by selectively overconsuming standard chow (a nutritionally balanced diet) rather than returning to the palatable glucose or lipid emulsion preferred at baseline. During clearance, feeding recovery consistently followed a strict macronutrient hierarchy: chow normalized and overshot first, followed by glucose, whereas lipids took the longest to recover. In effect, mice treated solid chow as a “safe” caloric source under semaglutide while actively avoiding liquid sugars and, to an even greater extent, fats. This pattern is well supported by evidence that GLP-1 receptor signaling selectively alters food choice rather than inducing a uniform anorexia. Preclinical choice paradigms have shown that systemic semaglutide and liraglutide markedly reduce the consumption of palatable candy, chocolate, and high-fat diets while preserving or increasing standard chow intake (Gabery et al., 2020; Gil Lievana et al., 2026; Jones et al., 2026; Raun et al., 2007), an effect mediated in part by mesolimbic GLP-1 receptors that selectively curb hedonic fat and sugar intake in favor of chow (Alhadeff et al., 2012; Dickson et al., 2012; Sandoval-Rodríguez et al., 2023; Zhu et al., 2025). Why, then, are lipids the last macronutrient to recover? Beyond central conditioned taste avoidance, where familiar chow is protected by latent inhibition relative to salient liquid tastants, peripheral digestive kinetics likely play a decisive role. Because dietary fats naturally delay gastric emptying more than carbohydrates, consuming liquid lipids during semaglutide clearance may exacerbate GLP-1-induced gastroparesis (Zhang et al., 2022). Indeed, both rodent studies on incretin agonists, tirzepatide (Geisler et al., 2023), and human clinical trials with semaglutide (Blundell et al., 2017; Gibbons et al., 2021) confirm a preferential and more persistent avoidance to high-fat foods compared with carbohydrates or balanced meals (Brindisi et al., 2019).

### Semaglutide induces taste avoidance rather than taste aversion, preserving taste palatability in mice

Nausea is the most frequent side effect of semaglutide in humans (Wilding et al., 2021), and in rats, which lack an emetic reflex, the highest dose used here has been shown to induce the intake of kaolin, a non-nutritive clay that rodents consume only when experiencing gastric distress, a model for pica (Ghidewon et al., 2022). Notably, we selected this high dose because it was also the most effective at inducing weight loss. Consequently, it is often assumed that semaglutide induces taste aversion; however, pica (a proxy for nausea) and taste aversion are two dissociable phenomena. Although the terms conditioned taste avoidance (measured by reduced intake in a bottle test) and conditioned taste aversion (measured by a negative shift on palatability) are often used interchangeably in the literature, taste avoidance is not always driven by taste aversion (Parker, 2003). Our data clearly show that semaglutide induces taste avoidance without affecting taste palatability; thus, despite inducing nausea, it is not sufficient to produce a frank taste aversion (**Figure 10**). Indeed, as noted above, our results align with recent reports demonstrating that chronic semaglutide preserves taste-driven palatability (lick-rates induced by sucrose) in brief-access tests in mice (Acosta et al., 2026) and even markedly increases intake of low concentrations of sucrose in rats (Cawthon et al., 2023). Likewise, in our lick-triggered taste alternation task, mice readily consumed glucose when it was passively delivered at the sipper (passive hits), and our Hidden Markov Model (HMM) confirmed that glucose-evoked oromotor responses remained intact under semaglutide, indicating that the drug did not alter hedonic ’liking.’ Thus, semaglutide primarily targets motivational and volitional processes.

### Limitations of the study

Several limitations should be noted. First, we examined only the acute effects of a single systemic dose of semaglutide. Because clinical use involves chronic administration, where sustained pharmacodynamics and receptor desensitization may occur (Son et al., 2026), future studies should test how chronic dosing (or drug-escalation) reshapes ingestive microstructure, taste categorization, and macronutrient recovery hierarchies. Second, our pharmacological evaluation was restricted to semaglutide. Whether other incretin-based therapeutics, such as the GLP-1/GIP dual agonist tirzepatide or the triple agonist retatrutide, produce the same effects on incentive motivation, taste categorization, and lipid avoidance remains to be determined, and will clarify whether these are class effects or ligand-specific phenotypes. Third, although our behavioral data show that semaglutide selectively impairs incentive motivation (wanting) while preserving glucose palatability and task rules, we did not record electrophysiological or neurochemical correlates. Given that the selective suppression of active foraging vigor and the dissociation between palatability and effort-related choice implicate mesolimbic circuitry (Zhu et al., 2025), future work should record dopamine dynamics (e.g.,accumbal or striatal dopamine release) during lick-triggered taste alternation tasks to link incretin receptor signaling with the dopaminergic encoding of glucose caloric value.

## Conclusion

From fueling mitochondrial energy production to driving the behavioral strategies mice employ to forage and feed, glucose serves as a fundamental energy source bridging cellular biochemistry and whole-organism behavior. Indeed, glucose is one of the most potent rewarding stimuli in nature (Sclafani et al., 2015), not by addiction but by evolution. The physiological need for this sugar has driven the evolution of sophisticated foraging strategies that continue to shape mammalian dietary habits (Gutierrez et al., 2020). Here, we comprehensively characterized glucose ingestive behavior under naturalistic conditions using the Homegustometer, revealing that glucose possesses a post-oral value that the artificial sweetener saccharin cannot match, rendering the search for truly iso-sweet or iso-preferred concentration pairs an illusion. More importantly, our novel lick-triggered taste alternation task revealed that semaglutide, with surgical precision, quiets ’food noise’ in mice by selectively impairing ’wanting,’ but not ’liking,’ for glucose.

## Acknowledgement

We thank Víctor Manuel García Gomez and Fabiola Olvera Hernández for the invaluable help with animal care, and Gustavo Hernández for building a prototype of the homegustometer. This project was supported by the SECIHTI Ciencia de Frontera CF-2023-G-518 to RG and PAPIIT IF102426 to ACR

## Funding

Secretaría de Ciencia, Humanidades, Tecnología e Innovación (CF-2023-G-518)

- Ranier Gutierrez

**PAPIIT IF102426**

- Alberto Caballero-Ruiz

## Supplemental Figures

**Figure 1–figure supplement 1.**
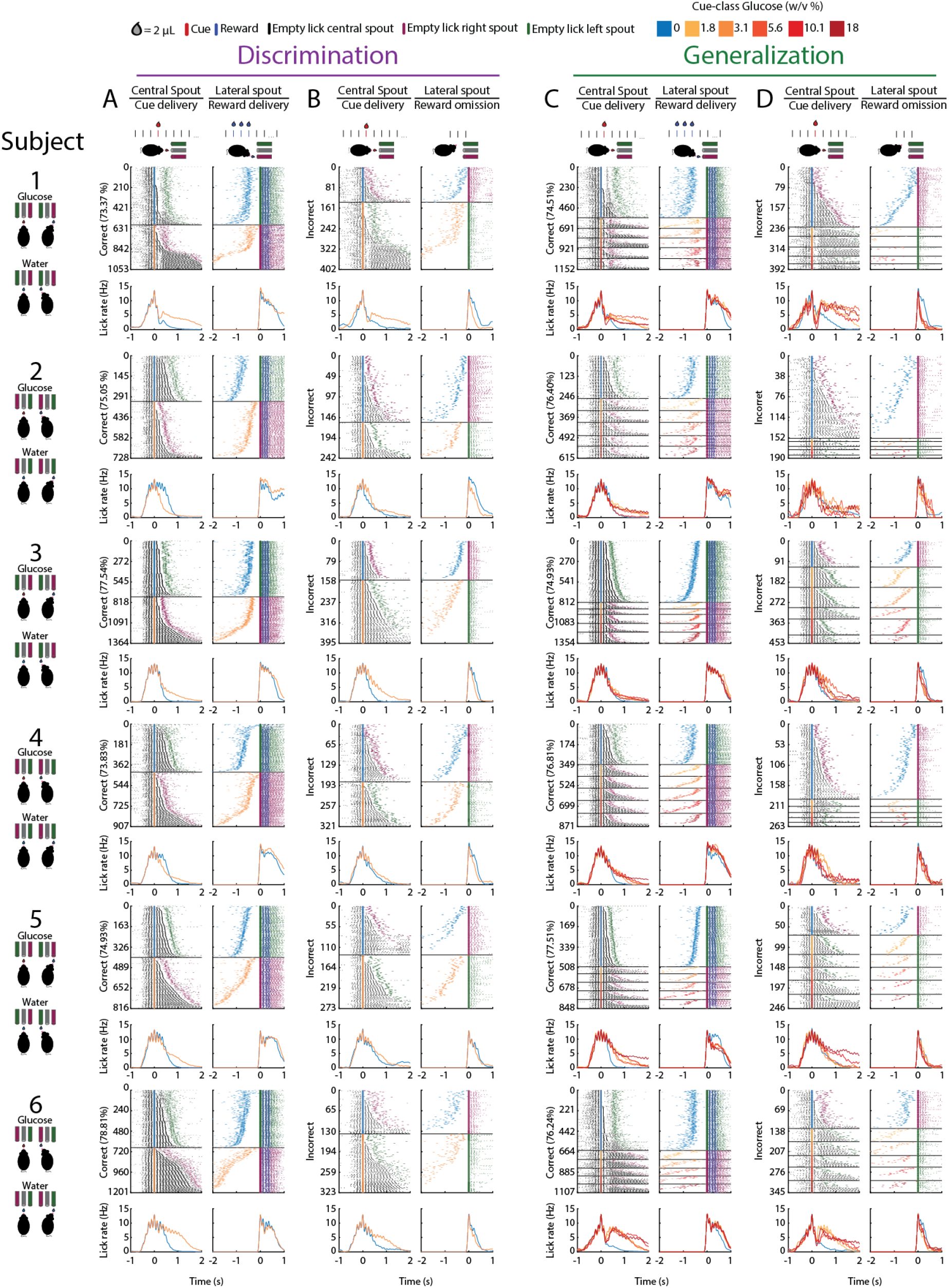
Individual licking profiles of all six subjects during criterion sessions of the discrimination and generalization phases. Behavioral profiles for each subject (1–6) across the three consecutive sessions in which the learning criterion (≥70% correct) was achieved. (**A, B**) Discrimination and (**C, D**) generalization sessions are separated into correct (**A, C**) and incorrect (**B, D**) trials. For each subject, upper panels show licking raster plots sorted vertically by lick bout duration, and lower panels display the corresponding mean peristimulus time histograms (PSTHs). Left columns are aligned to cue delivery at the central spout (t = 0 s), for visualization purposes only; licks in the central spout are plotted; right columns are aligned to the outcome at the lateral spouts (t = 0 s), only licks in the lateral sippers are shown plus the tick for cue delivery in the central spout: reward delivery on correct trials (**A, C**) and reward omission on incorrect trials (**B, D**). Cue ticks and PSTH traces are color-coded from blue to dark red according to cue-class glucose concentration (0, 1.8, 3.1, 5.6, 10.1, and 18% w/v); blue ticks at the lateral spouts indicate water reward deliveries. Black, magenta, and green ticks represent empty (dry) licks at the central, right, and left spouts, respectively. Schematics on the far left depict the stimulus–spout contingency for each subject; note that the glucose-associated lateral spout was counterbalanced across mice. Each drop delivered 2 μL.

**Figure 1—figure supplement 2.**
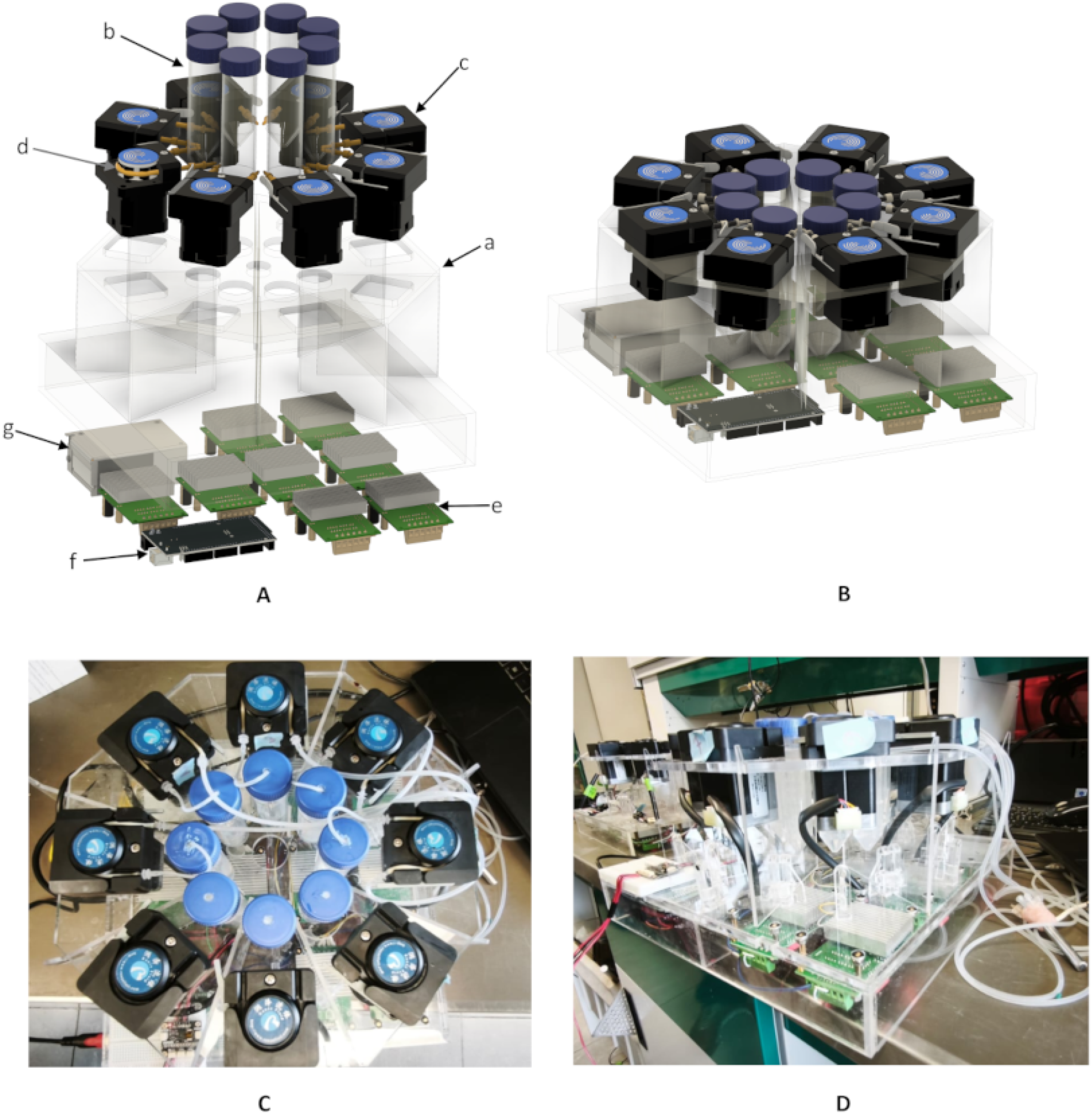
The Homegustometer dispenser system. As is well known in neuroscience, a gustometer is a specialized scientific device that delivers precise volumes of tastants to animals, triggered by lickometers, allowing researchers to measure and study behavioral and neural responses (Brosvic and Slotnick, 1986; Spector et al., 1990; Von Békésy, 1965). In rodents (especially mice), delivery volumes can be as small as a few microliters. Over time, gustometer dispenser designs have evolved from basic mechanical pumps to automated systems. A previous system from our laboratory could independently deliver multiple solutions using air-pressurized chromatography columns, with delivery controlled via solenoid valves (Fonseca et al., 2018; Gutierrez et al., 2010). However, these systems are limited to experiments lasting only a few hours and are unsuitable for long-term studies because volume-delivery accuracy is compromised by the changing fluid height in the chromatography column, which gradually reduces the dispensed volume over time. Here, we improve our system to run day and night continuously. The homegustometer dispenser system employed in this work can deliver up to eight different tastants and relies on peristaltic pumps to ensure dispense-volume accuracy independent of solution height. **Panel A** shows an exploded view of the proposed design. The main structure (a) has two levels: the upper level is octagonal and houses, near its center, eight chromatography columns (b) with a maximum capacity of 100 mL each. Surrounding these are eight stepper motor-driven peristaltic pumps (c) (model BJ-RZ1030, eight-roller mini-pump) equipped with Pharmed BPT-0064-008 tubing (d) (0.64 mm inner diameter, 2.64 mm outer diameter), both sourced from Nanjing Runze Fluid Control Equipment Co., Ltd. The lower level is square and houses the peristaltic pump controllers (TB6560 STEPPER MOTOR DRIVER) (e), which are managed by an Arduino MEGA module (f). The complete system is powered by a 12V–5A power supply (g). **Panel B** shows the system assembly, **Panel C** shows the top view of the assembled system, and **Panel D** shows the side view. The peristaltic pump setup allows for a maximum flow rate of 9 mL/min. Each pump is activated by contact lickometers (Med Associates Inc., VT, USA). Delivered tastant volumes range from 1 to 2 µL, which must be calibrated once to correspond to a specific number of stepper motor steps. Each tastant must be calibrated independently because the dispensed volume depends on the specific properties of the solution. The system error is less than 5%, and the delivery time is less than 5 milliseconds.

**Figure 2–figure supplement 1.**
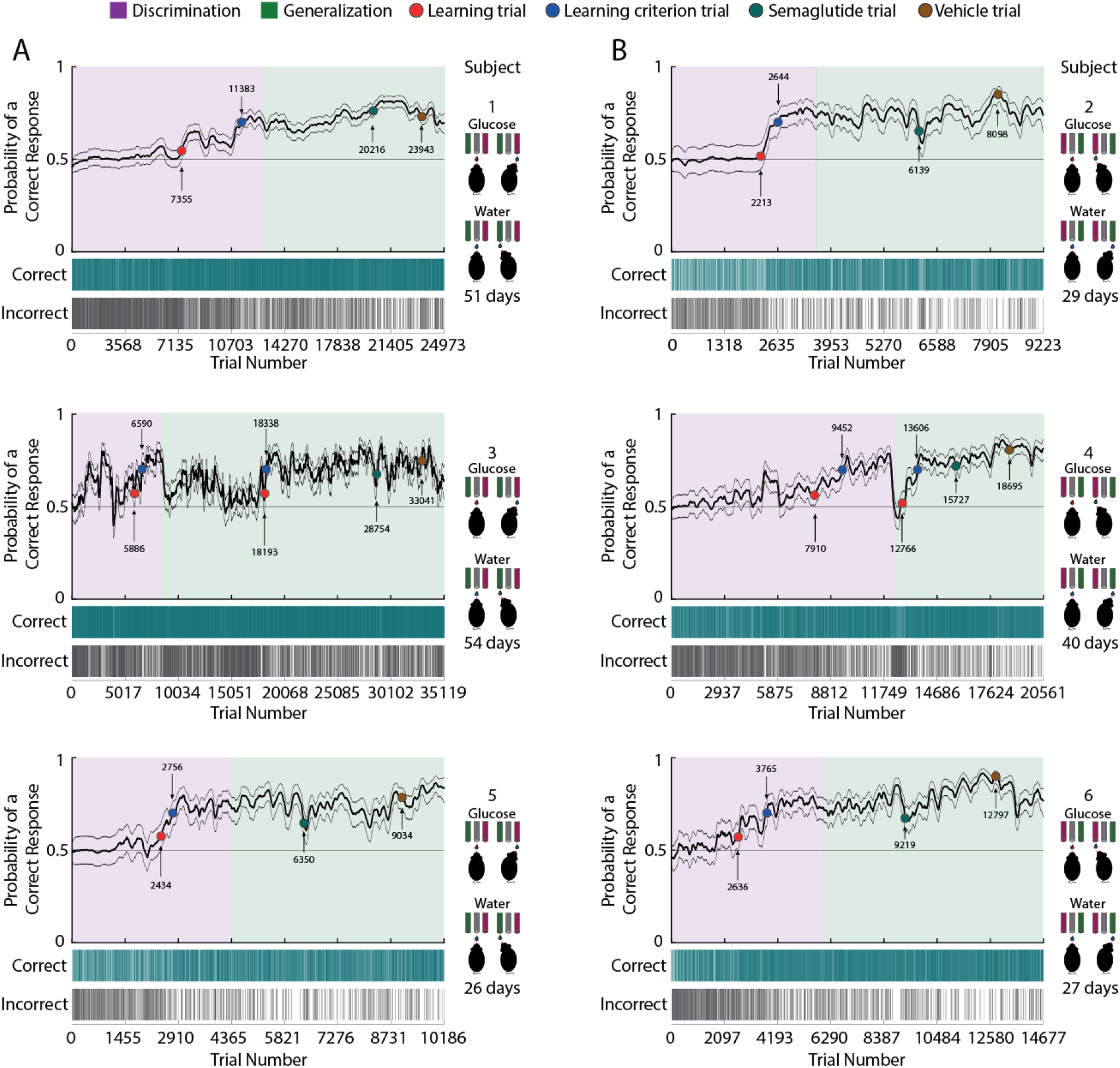
Individual trial-by-trial learning trajectories in the water–glucose categorization task. (A, **B)** State-space learning curves (upper) and trial-by-trial correct/incorrect choices (lower) for each of the six subjects, together with the total experimental duration in days (range: 26–54 days). The state-space model estimates the probability of a correct response (thick black line) with its 90% confidence intervals (thin black lines) at single-trial resolution. Background shading indicates trials performed during discrimination (purple) and generalization (green) sessions. The horizontal black line marks the chance level for this design (0.5). The learning trial (red dot) is the trial from which the lower confidence bound consistently exceeded chance, whereas the learning criterion trial (blue dot) is the first trial at which the estimated probability of a correct response reached 0.7. Dark teal and brown dots mark the first trial performed after semaglutide and vehicle administration, respectively; trial numbers for all annotated events are indicated. Schematics to the right of each panel depict the stimulus–spout contingency for each subject. Note that despite widely varying learning rates (learning criterion trial reached between ∼2,400 and ∼18,000 trials), all mice acquired the task, sustained performance above criterion during generalization, and showed the transient semaglutide-induced drop at the individual level.

**Figure 5-figure supplement 1.**
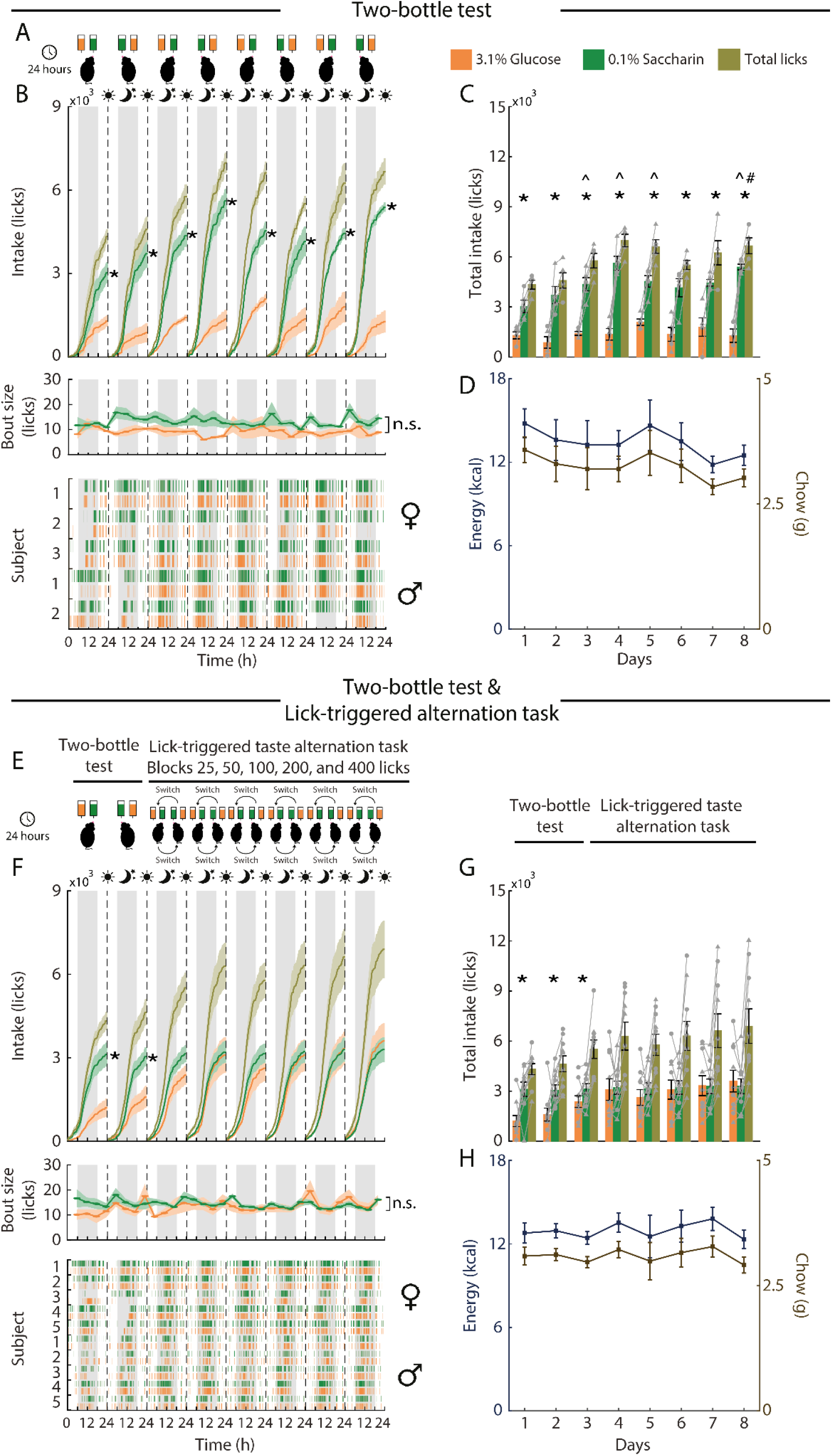
0.1% Saccharin over 3.1% glucose preference is mediated through sweeteners intensity and spatial sipper association. Two independent rodent cohorts were tested to assess preference between 0.1% saccharin and 3.1% glucose through 8 consecutive days. (A). Male (n = 2) and female mice (n = 3) were exposed to both caloric- and non-caloric sweetener in a prototypical two-bottle test. (E). The second group (n = 5 for both female and male) was evaluated for two days in conventional two-bottle tests, later they performed the lick-triggered taste alternation task swapping solution every 25, 50, 100, 200, and 400 licks for six consecutive days. Tastants were counterbalanced across days and subjects every 24 hours in all sessions in two-bottle tests. (B, F). Depicts cumulative licks (upper) across day (white and sun) and night (gray and moon) for 0.1% saccharin (green), 3.1% glucose (orange), and total licks (olive green). Vertical dashed lines indicate session boundaries. Asterisks denote statistical difference for pair tastants the same day (two-tailed K-S test, all *p*’s < 0.05). Indicates bout size, measured as count licks within a bout binned every six hours, for glucose and saccharin (middle). Licking rasters sorted by sex (♀ upper, ♂ lower). It is highlighted that the majority licking choices for saccharin were made in prototypical two-bottle tests (B) while intake in lick-triggered taste alternation task (F) mice did not exhibit a concrete non-caloric over caloric sweetener preference. (C, H). Total consumption per session across experimental days for 0.1% saccharin (green), 3.1% glucose (orange) and total licks (olive green) for two-bottle test (C) and two-bottle test and lick-triggered taste alternation task (G) groups. Note that irrespective of the behavioral paradigm, rodents effectuated the same amounts of total licks (olive green) basically differing how to distribute the solution choices. (D, H). Represent energy (left y-axis and blue) and chow (right y-axis and brown) intake across experimental days for two-bottle test (D) and two-bottle test and lick-triggered taste alternation task (H) mice. independently of the behavioral paradigm, subjects did not alter caloric demands. Data are plotted as mean ± SEM. Individual animals are represented by triangles and squares for females and males, respectively. Asterisks denote differences between solutions within a day. Carets and hashes represent differences between saccharin and glucose across experimental groups, respectively.

**Figure 7-figure supplement 1.**
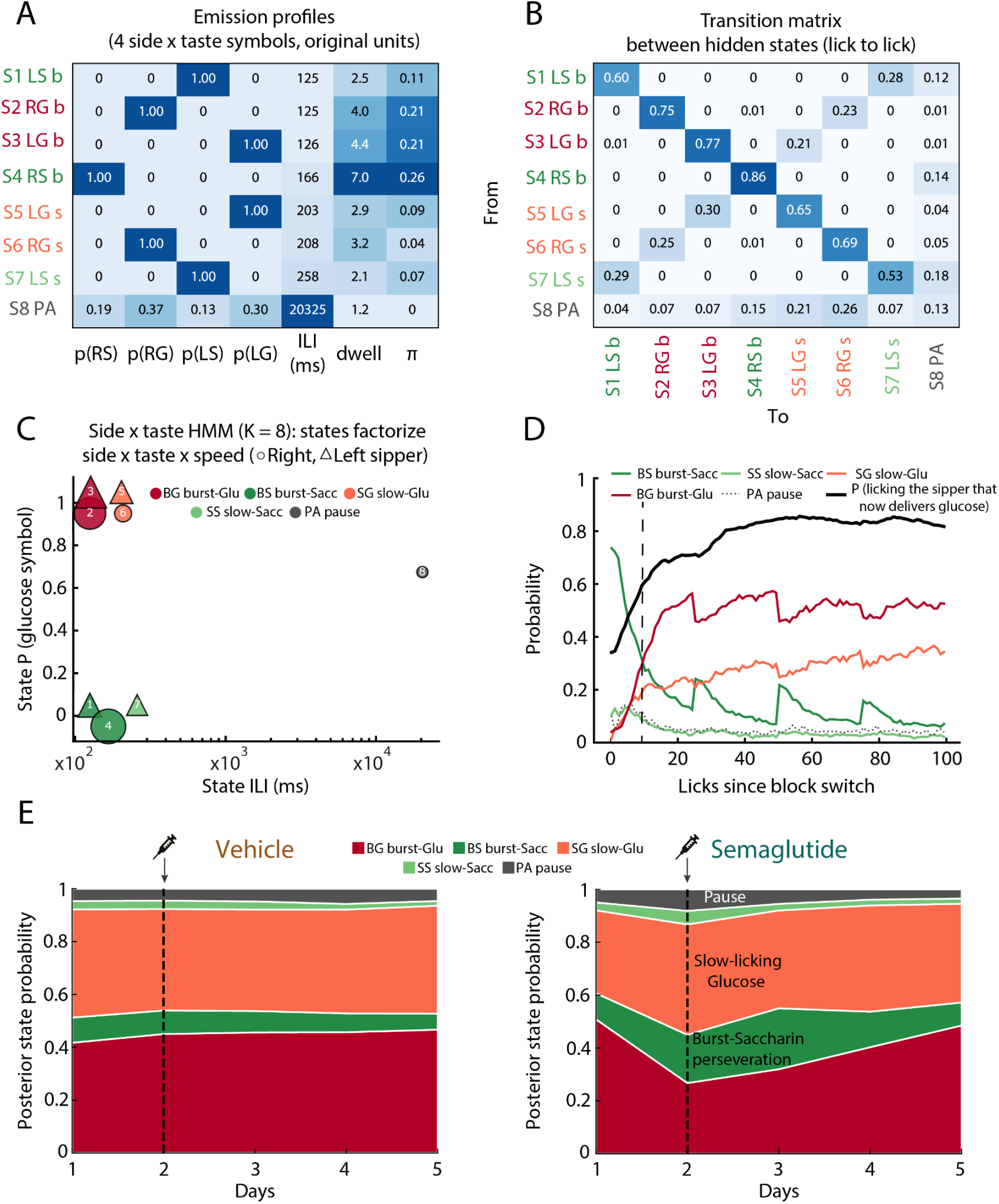
A side × taste Hidden Markov Model recovers a symmetric, pause-gated task architecture. A single Hidden Markov Model (HMM; K = 8 states) was fitted to the lick sequences of all 24 mice (n = 12 per group). Each lick was coded by its side × taste symbol (RS, right sipper–saccharin; RG, right sipper–glucose; LS, left sipper–saccharin; LG, left sipper–glucose) and its inter-lick interval (ILI). Emissions were modeled as the product of a categorical distribution over the four symbols and a Gaussian over log ILI. States are labeled by number, symbol and regime (b, burst; s, slow; PA, pause). **(A)** Emission profile of each state: symbol probabilities, mean ILI (ms, back-transformed from the log scale), expected dwell (licks) and initial probability (π). All seven active states are symbol-pure. The pause (S8) is the only state that emits all four symbols. **(B)** Lick-to-lick transition matrix (rows, origin state; columns, destination state). Off-diagonal transitions link the burst and slow states of the same symbol. No transitions connect the two sides directly, and only the pause row feeds both sippers. **(C)** States in emission space: mean ILI (log scale) versus P(glucose symbol). ○, right sipper; △, left sipper; marker area is proportional to π. The pause is the only mixed state (P(glucose) = 0.67). **(D)** Posterior probability of the five latent classes aligned to block switches (baseline). Each class pools the left and right states: BG, burst-glucose (S2, S3); BS, burst-saccharin (S1, S4); SG, slow-glucose (S5, S6); SS, slow-saccharin (S7); PA, pause (S8). The black line is the probability of licking at the sipper that now delivers glucose, the behavioral read-out of the reversal: the sippers never move, the tastes swap sides and the mouse must relocate. **(E)** Daily posterior (forward–backward gamma) class occupancy (forward–backward algorithm; mean across mice within each group) for vehicle (left) and semaglutide (right). Dashed line, injection at the start of day 2.

